# The cytosolic *Arabidopsis thaliana* cysteine desulfurase ABA3 delivers sulfur to the sulfurtransferase STR18

**DOI:** 10.1101/2020.03.25.008375

**Authors:** Benjamin Selles, Anna Moseler, Damien Caubrière, Sheng-Kai Sun, Morgane Ziesel, Tiphaine Dhalleine, Mathilde Hériché, Markus Wirtz, Nicolas Rouhier, Jérémy Couturier

**Author notes:** Université de Lorraine, CNRS, IMoPA, F-54000 Nancy, France. Institute of Crop Science and Resource Conservation (INRES)-Chemical Signalling, University of Bonn, 53113 Bonn, Germany (Anna Moseler). Agroécologie, AgroSup Dijon, CNRS, Université de Bourgogne, INRAE, Université de Bourgogne Franche-Comté, F-21000 Dijon, France. These authors contributed equally to this work (Benjamin Selles, Anna Moseler, Damien Caubrière). Corresponding author: Jérémy Couturier.

## Abstract

The biosynthesis of many sulfur-containing molecules depends on cysteine as a sulfur source. Cysteine desulfurase (CD) and rhodanese (Rhd) domain-containing protein families participate in the trafficking of sulfur for various metabolic pathways in bacteria and human, but their connection is not yet described in plants. The existence of natural chimeric proteins, however, containing both CD and Rhd domains in specific bacterial genera suggests a general interaction between both proteins. We report here the biochemical relationships between two cytosolic proteins from *Arabidopsis thaliana*, a Rhd domain containing protein, the sulfurtransferase 18 (STR18), and a CD isoform referred to as ABA3, and compare these biochemical features to those of a natural CD-Rhd fusion protein from the bacterium *Pseudorhodoferax sp*.. We observed that the bacterial enzyme is bifunctional exhibiting both CD and STR activities using L-cysteine and thiosulfate as sulfur donors but preferentially uses L-cysteine to catalyze trans-persulfidation reactions. *In vitro* activity assays and mass spectrometry analyses revealed that STR18 stimulates the CD activity of ABA3 by reducing the intermediate persulfide on its catalytic cysteine thereby accelerating the overall transfer reaction. Both proteins interact *in planta* and form an efficient sulfur relay system whereby STR18 catalyzes trans-persulfidation reactions from ABA3 to the model acceptor protein roGFP2. In conclusion, the ABA3-STR18 couple likely represents an uncharacterized pathway of sulfur trafficking in the cytosol of plant cells, independent of ABA3 function in molybdenum cofactor maturation.

---

Sulfur is an essential macronutrient playing pivotal roles in the physiology and development of all organisms as it is present in two amino acids, cysteine and methionine, but also in many other molecules such as sulfolipids, thionucleosides, vitamins (thiamin, biotin, lipoic acid) and iron-sulfur (Fe-S) clusters or molybdenum cofactors (Moco) (1). In plants, cysteine is synthesized *via* the reductive assimilation of sulfate, and is the source of reduced sulfur for the biosynthesis of most sulfur-containing cofactors or molecules mentioned above (1). The common feature in the biosynthetic schemes involving the formation of sulfur-containing compounds in bacteria and eukaryotes is the expression of specific proteins that activate the sulfur from cysteine and transfer it to target acceptor proteins. Then, the nature of sulfur acceptors and their chemical functionality dictate the direction and the flow of sulfur transfer. Cysteine desulfurases (CDs) constitute a family of enzymes responsible for the sulfur transfer from cysteine to acceptor molecules (1). These ubiquitous proteins are pyridoxal 5’-phosphate (PLP)-dependent enzymes catalyzing the desulfuration of cysteine, leading to the formation of a persulfide group on a catalytic cysteine and the concomitant release of alanine (2). Then, the efficiency and specificity of sulfur transfer to acceptor molecules vary according to the subclass of CDs and the type of sulfur acceptors.

The NifS protein from *Azotobacter vinelandii* was the first CD characterized for its involvement in the maturation of the Fe-S cluster present in nitrogenase (3). This functional assignment led to the subsequent identification of the IscS paralog, that serves as a general system for the maturation of other Fe-S proteins but also for providing sulfur present in other molecules (4). In addition to IscS, *Escherichia coli* possesses two other CD isoforms, namely SufS and CsdA (5, 6). Bacterial and eukaryotic CDs share a similar fold and assemble as dimers but two groups have been distinguished based on distinct structural differences and reactivities (6). IscS- and NifS-like proteins are members of group I. They contain a 12-residue insertion in an exposed loop containing the catalytic cysteine. In *Ec*IscS, this extension is sufficiently flexible to allow the direct transfer of sulfur to multiple biological partners (7). SufS- and CsdA-like proteins belong to the group II and the loop containing the catalytic cysteine is shorter (8, 9). For this reason, they form a two-component system with specific activators/sulfur acceptors, *i*.*e. Ec*SufE and *Ec*CsdE with *Ec*SufS and *Ec*CsdA respectively, or SufU with SufS in *Bacillus subtilis* (10–13).

In plants, mitochondrial NFS1 and plastidial NFS2 are the CDs providing the sulfur required for Fe-S cluster assembly in both mitochondria and cytosol or in chloroplasts, respectively (14, 15). The CD activity of NFS2 is relatively low in the absence of the specific SUFE1-3 activators (16–18). A third CD isoform, ABA3, is localized in the cytosol of plants and involved in Moco sulfuration, thus participating in the activation of aldehyde oxidase (AO) and xanthine dehydrogenase (XDH). These Moco-containing enzymes are involved in abscisic acid biosynthesis and in purine degradation respectively (19, 20). ABA3 is formed by two domains, an N-terminal aminotransferase class V domain (InterPro: IPR000192) responsible for the CD activity, as in NFS1 and NFS2. In addition, the protein possesses a C-terminal MOSC domain (InterPro: IPR005302 and IPR005303) responsible for the final incorporation of a sulfur atom into the Moco precursor (20, 22).

Similar trans-persulfidation reactions between CDs and other sulfur carrier proteins occur during the biosynthesis of sulfur-containing molecules. Among these sulfur carrier proteins are sulfurtransferases (STRs), widespread enzymes present in bacteria, archaea and eukarya. They possess a characteristic rhodanese (Rhd) domain usually containing a conserved catalytic cysteine present in a specific Cys-X-X-Gly-X-Arg signature (23, 24). This cysteine is mandatory to the catalytic activity of STRs since a cysteine persulfide intermediate is formed during trans-persulfidation reactions. Three different STR classes have been defined with respect to their modular organizations and substrate specificities (23–25). STRs with a single Rhd domain use preferentially thiosulfate as a sulfur donor *in vitro* and are referred to as thiosulfate-STRs (TSTs, InterPro: IPR001307). Those possessing two Rhd domains use preferentially 3-mercaptopyruvate (3-MP) as a sulfur donor *in vitro* and were named 3-mercaptopyruvate (3-MP)-STRs (MSTs, InterPro: IPR036873). Mammals possess an additional STR isoform with two Rhd domains, named Rhobov, that uses sulfite and glutathione persulfide to synthesize thiosulfate. Additional STR proteins contain one Rhd domain fused to one or several protein domains with another function conferring them specific roles (23, 24).

Examples of interaction between CDs and STRs in non-plant organisms have suggested a hub function for CD/STR couples as sulfide/sulfur moieties are required for various metabolic pathways. In *E. coli*, the sulfur transfer from IscS to the STRs, ThiI or YnjE, is involved in thiamine biosynthesis or in tRNA thiolation and Moco biosynthesis respectively (26, 27). A similar sulfur relay system exists in the cytosol of yeast and human. The human STR isoform, TUM1, participates in the biosynthesis of Moco cofactor, receiving sulfur from NFS1 (28). Moreover, TUM1 proteins ensure sulfur transfer to another STR isoform, referred to as Uba4 in yeast or MOCS3 in human, playing a role in tRNA thiolation (29).

Such sulfur transfer relays should be universal considering the existence of natural chimeric proteins containing both CD and Rhd domains in specific bacterial genera. However, their properties have not been characterized, nor the existence of a comparable system in plants. Previous studies on plant CDs were mostly focused on their role in Fe-S cluster biogenesis (NFS1, NFS2) and Moco sulfuration (ABA3), not on a possible interaction with STRs (14, 15, 21). Hence, we have investigated the biochemical properties and interactions between the *Arabidopsis thaliana* cytosolic ABA3 and STR18 and compared these biochemical features to those of a natural CD-Rhd fusion protein present in the bacterium *Pseudorhodoferax sp*.. We demonstrated that the bacterial enzyme is bifunctional exhibiting both CD and thiosulfate-dependent STR activities. Using redox-sensitive GFP (roGFP2) as a model acceptor protein, we showed the ability of CD-Rhd to catalyze efficiently trans-persulfidation reaction from L-cysteine but not thiosulfate to roGFP2. Concerning plant proteins, *in vitro* activity assays and mass spectrometry analyses revealed that STR18 stimulates the CD activity of ABA3 by reducing the persulfide formed on the CD catalytic cysteine. Using roGFP2 assay, we demonstrated the ability of STR18 to catalyze trans-persulfidation reactions from ABA3 to roGFP2. Finally, split-luciferase complementation assays revealed that both proteins interact *in planta*. Considering all these data, our study reveals that the ABA3-STR18 couple likely represents a new pathway of sulfur trafficking in the cytosol of *A. thaliana*.

## RESULTS

### The natural CD-Rhd fusion protein of Pseudorhodoferax sp. is a bifunctional enzyme

Genomic analyses like gene clustering, gene co-occurrence or gene fusion are powerful tools to predict functional associations. For instance, the existence of natural fusions in some organisms often reflects a functional interaction in other organisms in which the constituting protein domains are expressed as separate proteins. By interrogating the STRING database (https://string-db.org/) using the COG1104 specific to CDs, we have noticed the existence of both adjacent CD and Rhd genes and natural CD-Rhd fusion genes/proteins in several bacteria. We focused our attention on a CD-Rhd isoform from *Pseudorhodoferax sp*. Leaf274. The presence of conserved catalytic cysteine residues in each protein domain suggests that this protein should possess both cysteine desulfurase and TST-type sulfurtransferase activities (Fig. 1A) (8, 23). The corresponding His-tagged recombinant protein exhibited a yellow color after purification. Additionally, the UV-visible absorption spectrum exhibited an absorption band at 418 nm characteristic for a bound-PLP cofactor as in characterized CDs (Fig. 1B) (9). Analytical gel filtration analysis demonstrated that CD-Rhd eluted predominantly in a peak corresponding to an apparent volume/molecular mass of 108 kDa (Fig. 1C). From the theoretical molecular mass of CD-Rhd (54 kDa), we concluded that this protein formed homodimers as observed for other CDs (7, 8).

**Figure 1.**
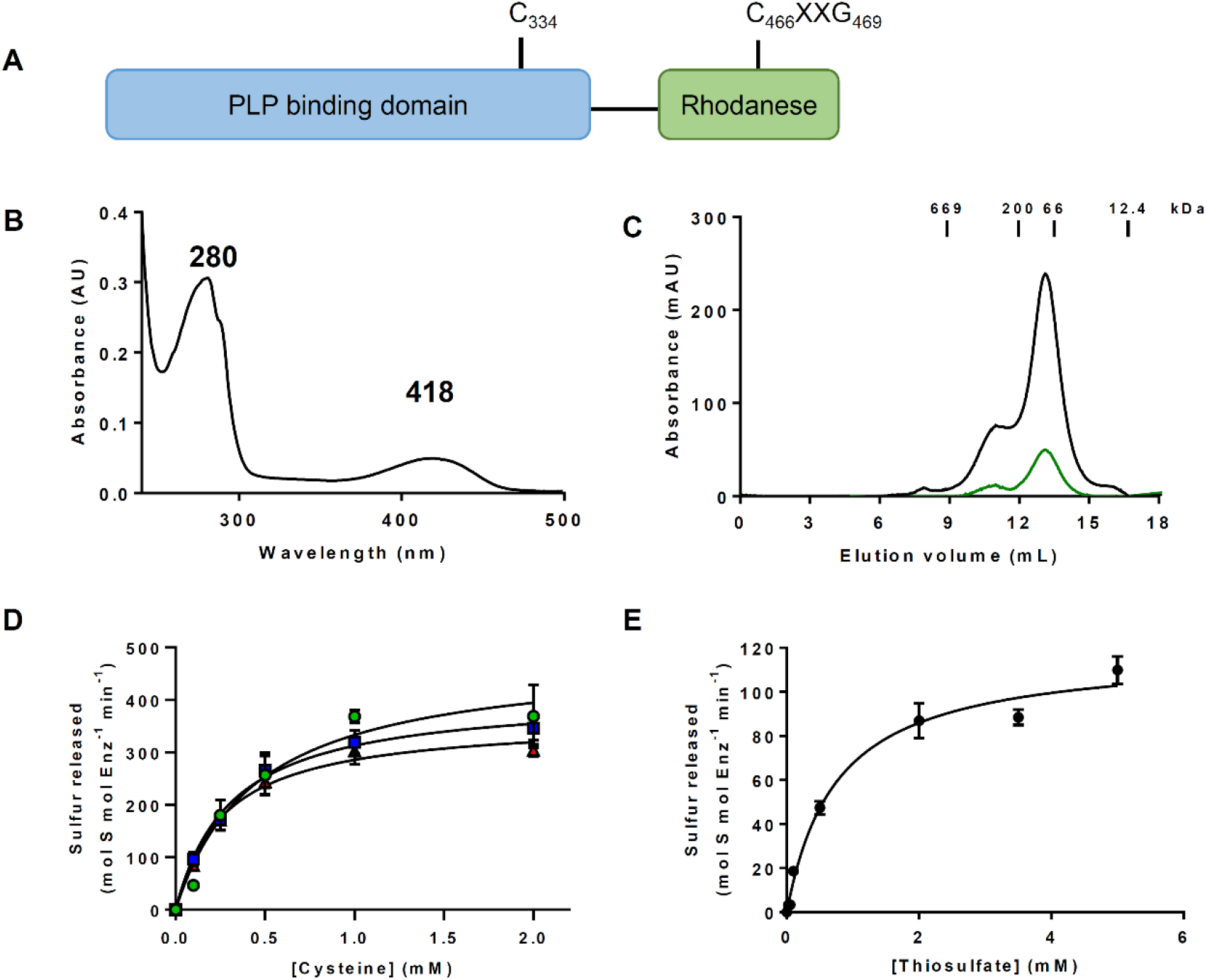
*Pseudorhodoferax* CD-Rhd fusion has a dual activity profile. A. Modular organization of the *Pseudorhodoferax* CD-Rhd fusion (WP_056898193.1) presenting the position of the presumed catalytic cysteines of both CD and Rhd domains. B. UV-visible absorption spectrum of the purified N-terminal His-tagged recombinant CD-Rhd recorded in a 30 mM Tris-HCl pH 8.0 buffer. C. Analytical gel filtration (Superdex S200 10/300 column, GE Healthcare) of His-tagged recombinant CD-Rhd (100 μg). The presence of the polypeptide and of the PLP cofactor have been detected by measuring the absorbance at 280 nm (dark line) and 418 nm (green line), respectively. The apparent molecular weight of CD-Rhd was estimated from the separation of the indicated standards. D. Steady-state kinetic parameters of the cysteine desulfurase activity. Reactions were performed in the presence of 10 nM CD-Rhd, increasing concentrations of L-cysteine (0 to 2 mM) and in the presence of various reductants, either 5 mM of DTT (blue squares), or 5 mM GSH (green circles) or 5 mM β-mercaptoethanol (red triangles). The data are represented as mean ± SD of three independent experiments. E. Steady-state kinetic parameters of the thiosulfate sulfurtransferase activity. Reactions were performed in the presence of 100 nM CD-Rhd, increasing concentrations of thiosulfate (0 to 5 mM) and 5 mM β-mercaptoethanol. The data are represented as mean ± SD of three independent experiments.

We have then evaluated the capability of the fusion protein to use L-cysteine or thiosulfate as substrates and determined the kinetic parameters of the reactions. The CD activity (*i*.*e*. cysteine desulfuration with the concomitant formation of a persulfide on catalytic cysteine) was monitored under steady-state conditions by measuring the release of H_2_S from the persulfidated protein in the presence of chemical or physiological reducing acceptors (Fig. 1D). Catalytic efficiencies (*k*_*cat*_/*K*_*M*_) of 2.2 × 10^4^ M^-1^ s^-1^, 1.8 × 10^4^ M^-1^ s^-1^ and 2.2 × 10^4^ M^-1^s^-1^ have been measured in the presence of DTT, reduced glutathione (GSH) and β-mercaptoethanol (β-ME), respectively (Table 1), thus validating the CD activity of the fusion. The activity of the Rhd domain was also evaluated by monitoring the release of H_2_S in the presence of β-ME but providing thiosulfate as the canonical substrate of TST-type STRs. The catalytic efficiency of the reaction was 2.7 × 10^3^ M^-1^ s^-1^ and the apparent *K*_*M*_ value for thiosulfate was 756 ± 51 μM (Fig. 1E, Table 1), thus validating the STR activity of the fusion. Our results demonstrate that the CD-Rhd chimeric protein from *Pseudorhodoferax* is bifunctional having dual *in vitro* activities using L-cysteine or thiosulfate as a sulfur donor.

**Table 1.** Kinetic parameters of cysteine desulfurase and thiosulfate sulfurtransferase activities of CD-Rhd and its C466S variant.

| Donor | Reductant | $K_M$ ($\mu\text{M}$) | $k_{cat}$ ($\text{s}^{-1}$) | $k_{cat}/K_M$ ($\text{M}^{-1} \text{s}^{-1}$) |
| --- | --- | --- | --- | --- |
| **CD-Rhd** |  |  |  |  |
| L-cysteine | DTT | $310 \pm 38$ | $6.8 \pm 0.5$ | $2.2 \pm 0.2 \times 10^4$ |
| L-cysteine | GSH | $476 \pm 146$ | $8.2 \pm 1.2$ | $1.8 \pm 0.2 \times 10^4$ |
| L-cysteine | β-ME | $273 \pm 26$ | $6.1 \pm 0.2$ | $2.2 \pm 0.1 \times 10^4$ |
| Thiosulfate | β-ME | $756 \pm 51$ | $2.0 \pm 0.1$ | $2.7 \pm 0.3 \times 10^3$ |
| **CD-Rhd C466S** |  |  |  |  |
| L-cysteine | DTT | $31 \pm 5$ | $1.01 \pm 0.01$ | $3.3 \pm 0.6 \times 10^4$ |
| L-cysteine | GSH | $597 \pm 168$ | $0.93 \pm 0.12$ | $1.6 \pm 0.2 \times 10^3$ |
| L-cysteine | β-ME | $346 \pm 79$ | $0.53 \pm 0.03$ | $1.6 \pm 0.3 \times 10^3$ |
| Thiosulfate | β-ME | n.d. | n.d. | n.d. |

The apparent $K_M$ and turnover values ( $k_{cat}$ ) were calculated by non-linear regression using the Michaelis-Menten equation. The data are represented as mean ± SD of three independent experiments. n.d. = not detected.

### Rhd domain promotes CD activity and trans-persulfidation reactions

Although the CD-Rhd protein is bifunctional, its CD activity was ∼8-fold higher than its TST activity. The Rhd domain only possesses the characteristic catalytic cysteine (Cys466) and thus could represent a sulfur acceptor for the CD domain. To determine the importance of the cysteine in the Rhd domain for the recorded CD activity, we analyzed the biochemical properties of a CD-Rhd C466S variant (Fig. S1, Table 1). Similar to CD-Rhd, the His-tagged CD-Rhd C466S recombinant protein exhibited a UV-visible spectrum with two absorption bands at 280 nm and 418 nm (Fig. S1A) and existed as a dimeric form (Fig. S1B). The absence of TST activity confirmed that the cysteine of the Rhd domain is mandatory for this activity (Fig. S1C). Concerning CD activity, the CD-Rhd C466S variant is still active despite its catalytic efficiency was ∼10-fold lower in the presence of GSH or β-ME (1.6 × 10^3^ M^-1^ s^-1^) compared with the activity of CD-Rhd. This is notably explained by a decrease of the apparent *k*_*cat*_ value by a factor of 10 (Fig. S1D, Table 1). On the contrary, CD activity did not significantly vary in the presence of DTT because the decrease of the apparent *k*_*cat*_ by a factor of 6 is compensated by a change in the apparent *K*_*M*_ value for cysteine (Fig. S1D, Table 1). The decrease in the turnover number of CD-Rhd C466S suggests that the Rhd domain stimulates the CD activity of the fusion with the catalytic cysteine probably serving as a persulfide-relay system.

To study the potential sulfur relay role of the Rhd domain, we investigated the capability of CD-Rhd and its C466S variant to transfer a sulfur atom to a protein substrate. In the absence of known CD-Rhd partners, we used roGFP2 that has been recently shown to act as an efficient sulfur acceptor for Rhd domain-containing proteins (30). We have first tested the oxidation of a pre-reduced roGFP2 in the presence CD-Rhd or its C466S variant and L-cysteine (Fig. 2A). L-cysteine alone had no effect on roGFP2 oxidation. On the contrary, the combination of L-cysteine with CD-Rhd led to an efficient roGFP2 oxidation (Fig. 2A). The reaction is much slower in the presence of the CD-Rhd C466S variant. These results demonstrated that a functional Rhd domain is necessary for an optimal reaction. The catalytic cysteine of the Rhd domain likely promotes the sulfur transfer from the catalytic cysteine of the CD domain to roGFP2 catalyzing a trans-persulfidation reaction between both proteins. Considering the TST activity of CD-Rhd, we performed similar experiments using thiosulfate instead of L-cysteine as a sulfur donor. Thiosulfate alone had not effect on roGFP2 and the CD-Rhd fusion was very poorly able to catalyze roGFP2 oxidation (Fig. 2B). No oxidation was observed with the CD-Rhd C466S variant (Fig. 2B). Altogether, these findings indicate that CD-Rhd preferentially uses L-cysteine as sulfur donor and the Rhd domain promotes trans-persulfidation reaction between CD domain and protein partner.

**Figure 2.**
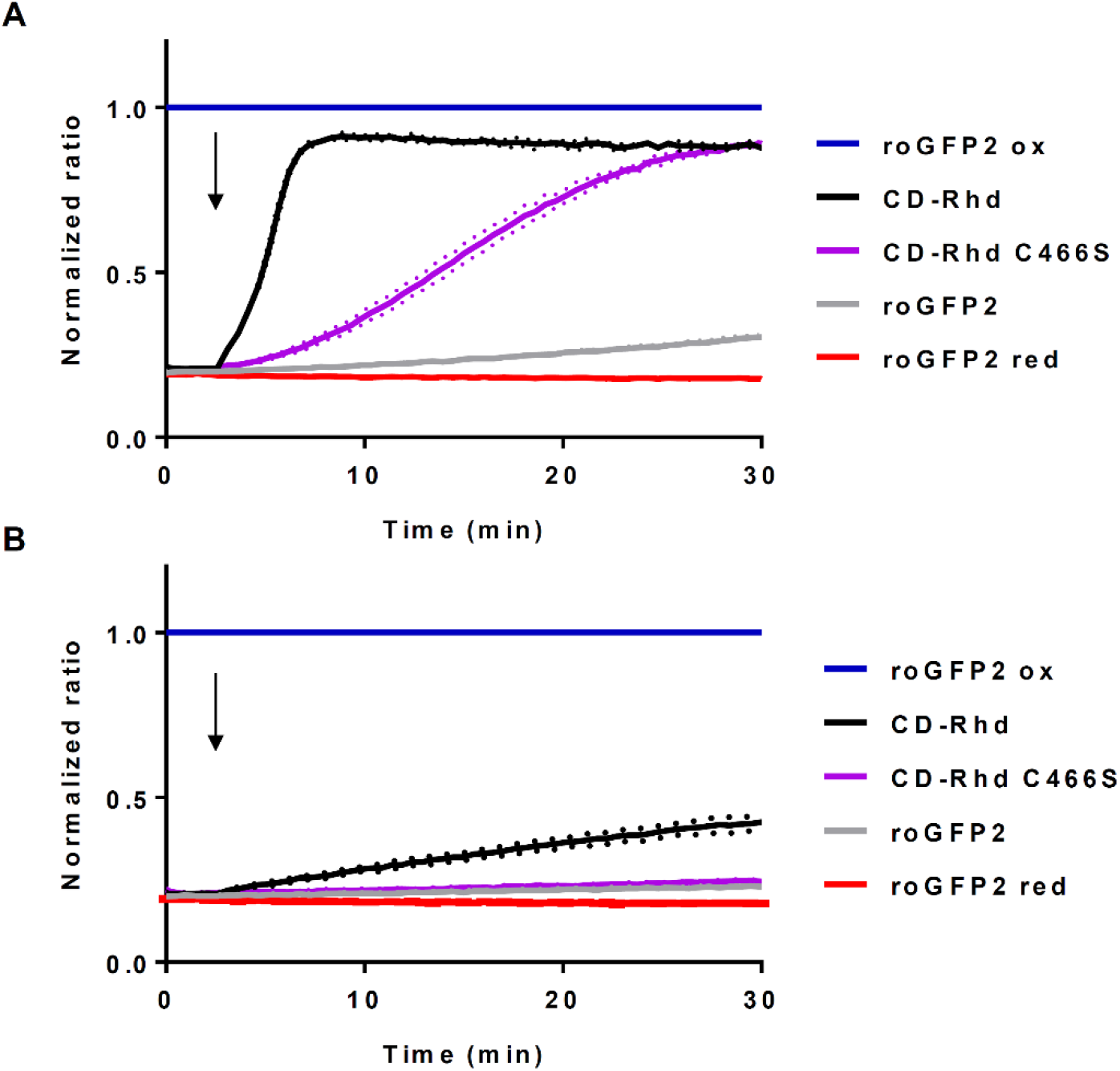
CD-Rhd catalyzes the oxidation of roGFP2 via trans-persulfidation. Persulfide-dependent oxidation kinetics of 1 μM roGFP2 in the presence of 5 μM CD-Rhd or the C466S variant and 1 mM L-cysteine (A) or 1 mM thiosulfate (B). Arrows indicate the addition of the respective substrate after 3 min. The fully reduced or oxidized roGFP2 used as references were obtained after incubation with 10 mM DTT or H_2_O_2_ respectively. The ratio of 400/480 nm was normalized to the respective value of maximal oxidation by H_2_O_2._ The data are represented as mean ± SD of three independent experiments.

### A. thaliana STR18 stimulates the cysteine desulfurase activity of ABA3

The results obtained with this chimeric protein prompted us to investigate the interaction between *A. thaliana* orthologs that exist as separate proteins, focusing on the cytosolic ABA3 and STR18, a single Rhd-domain containing protein (22, 24, 31, 32, 33). STR18 possesses two cysteine residues, Cys47 and Cys89, the latter corresponding to the catalytic cysteine present in the Cys-X-X-Gly-X-Arg signature typical of the Rhd domain (24, 32). TST activity assays were first performed with STR18 and both C47S and C89S variants. The STR18 C89S variant was inactive whereas the catalytic efficiency of the STR18 C47S variant was only marginally affected (Fig. S2, Table 2). This indicated that STR18 exhibits TST activity, that Cys89 is mandatory and Cys47 dispensable. The influence of STR18 and its variants on CD activity of ABA was then evaluated. The turnover for cysteine desulfuration of ABA3 was found to be ∼2-fold higher in the presence of STR18 (0.66 *vs* 1.39 mole sulfur mole enz^-1^ min^-1^), indicating a stimulating effect of STR18 on the CD activity of ABA3 (Fig. 3A). As expected, the STR18 C47S variant also stimulated the CD activity of ABA3 while the STR18 C89S variant did not (Fig. 3A).

**Table 2.** Kinetic parameters of thiosulfate sulfurtransferase activity of STR18, C47S and C89S variants.

| | $K_M$ ($\mu\text{M}$) | $k_{cat}$ ($\text{s}^{-1}$) | $k_{cat}/K_M$ ($\text{M}^{-1} \text{s}^{-1}$) |
| --- | --- | --- | --- |
| STR18 | $527 \pm 8$ | $5.1 \pm 0.1$ | $9.7 \pm 0.2 \times 10^3$ |
| STR18 C47S | $333 \pm 15$ | $2.5 \pm 0.1$ | $7.5 \pm 0.2 \times 10^3$ |
| STR18 C89S | n.d. | n.d. | n.d. |

The TST activity of STR18, C47S and C89S variants was monitored in the presence of varying concentrations of thiosulfate as described in the “Experimental procedures” section. The apparent $K_M$ and turnover values ( $k_{cat}$ ) were calculated by non-linear regression using the Michaelis-Menten equation. The data are represented as mean ± SD of three independent experiments. n.d. stands for not detected.

**Figure 3.**
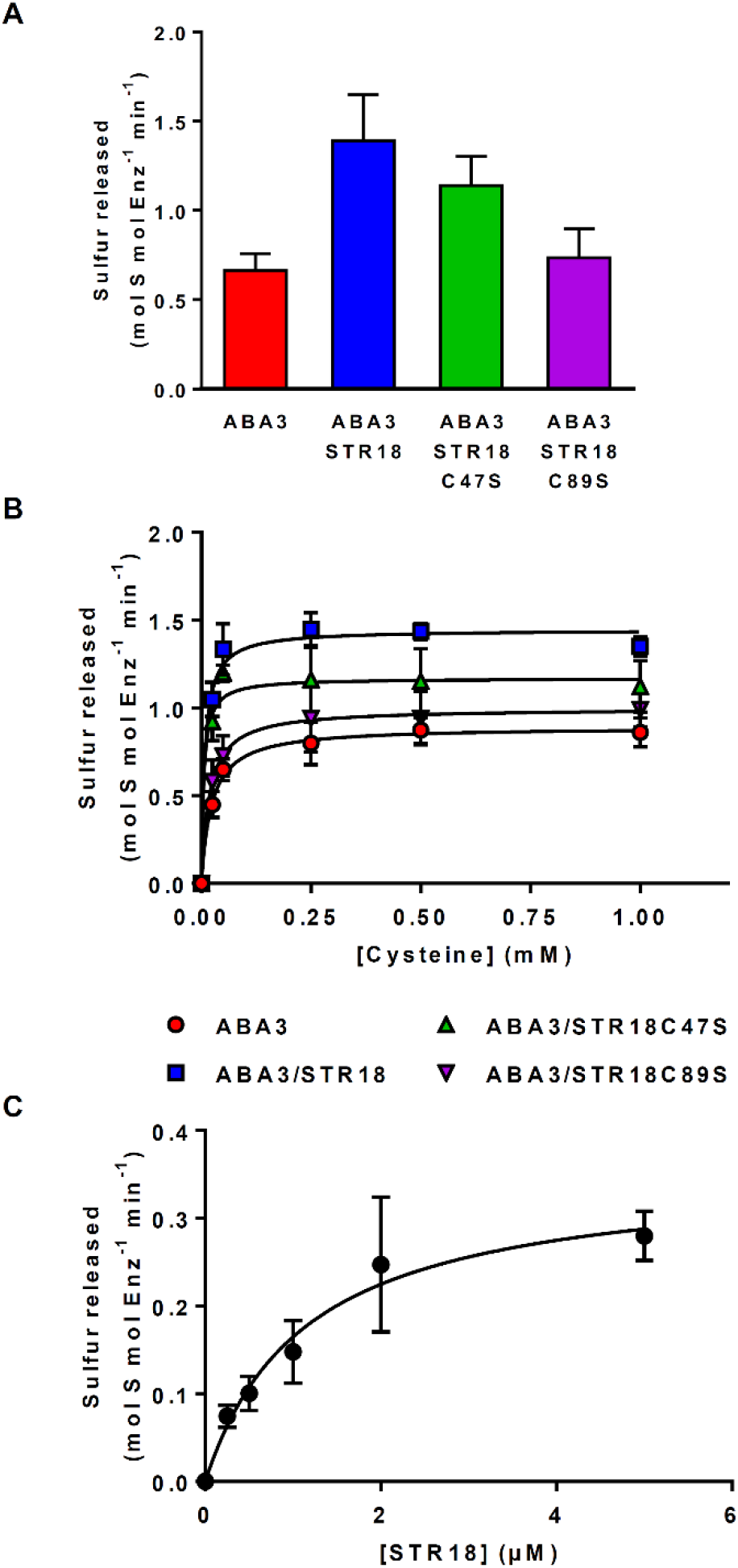
STR18 stimulates ABA3 activity. A. Effect of STR18 on ABA3 cysteine desulfurase activity. CD activity was measured using 1 μM ABA3, 500 μM L-cysteine and 1 mM DTT in the presence or absence of 5 μM STR18. B. Steady-state kinetic parameters of the CD activity of ABA3 alone or in the presence of STR18 or its cysteinic variants. CD activity was measured in the presence of 1 μM ABA3, increasing concentrations of L-cysteine (0 to 1 mM), 5 mM DTT and 5 μM STR18 when present. C. Steady-state kinetic parameters of the CD activity of ABA3 in the presence of STR18. Activity was measured in the presence of 0.5 μM ABA3, 500 μM L-cysteine, 5 mM DTT and increasing concentrations of STR18 (0 to 5 μM). The data are represented as mean ± SD of three independent experiments.

These results prompted us to investigate the interaction of ABA3 with STR18 and its monocysteinic variants and to determine the steady-state kinetic parameters associated with CD activity (Fig. 3B, Table 3). In the absence of STR18, the apparent *K*_*M*_ values of ABA3 for L-cysteine was 23 ± 6 μM and the deduced catalytic efficiency was 680 M^-1^ s^-1^ (Fig. 3B, Table 3). In the presence of STR18, ABA3 was 5-fold more efficient and this is notably explained by a decrease of the apparent *K*_*M*_ value for L-cysteine by a factor of 3 (Fig. 3B, Table 3). Similar kinetic parameters were obtained using the STR18 C47S variant, while, as expected, the CD activity of ABA3 was not stimulated in the presence of the STR18 C89S variant as illustrated by the apparent *K*_*M*_ value for L-cysteine and the *k*_*cat*_/*K*_*M*_ value close to those determined for ABA3 alone (Fig. 3B, Table 3).

**Table 3.** Kinetic parameters of cysteine desulfurase activity of ABA3.

| | $K_M$ ($\mu\text{M}$) | $k_{cat}$ ($\text{s}^{-1}$) | $k_{cat}/K_M$ ($\text{M}^{-1} \text{s}^{-1}$) |
| --- | --- | --- | --- |
| L-cysteine |  |  |  |
| ABA3 | $23 \pm 6$ | $0.015 \pm 0.001$ | $6.8 \pm 1.6 \times 10^2$ |
| ABA3 + STR18 | $8 \pm 1$ | $0.024 \pm 0.001$ | $3.2 \pm 0.5 \times 10^3$ |
| ABA3 + STR18 C47S | $6 \pm 1$ | $0.019 \pm 0.002$ | $3.1 \pm 0.4 \times 10^3$ |
| ABA3 + STR18 C89S | $19 \pm 3$ | $0.017 \pm 0.002$ | $9.3 \pm 2.2 \times 10^2$ |
| STR18 |  |  |  |
| ABA3 | $1.2 \pm 0.2$ | $0.006 \pm 0.001$ | $5.1 \pm 0.8 \times 10^3$ |

The CD activity of ABA3 was monitored in the presence of varying concentrations of L-cysteine and with or without STR18 as described in the “Experimental procedures” section. The apparent $K_M$ and turnover values ( $k_{cat}$ ) were calculated by non-linear regression using the Michaelis-Menten equation. The data are represented as mean ± SD of three independent experiments.

Altogether, these data indicate that STR18 stimulates the CD activity of ABA3 by increasing ABA3 affinity for L-cysteine. To further characterize ABA3-STR18 interaction, the CD activity of ABA3 was monitored in the presence of 500 μM L-cysteine and of increasing STR18 concentrations. This allowed us to determine apparent *K*_*M*_ values of ABA3 for STR18 of 1.2 ± 0.2 μM (Fig. 3C, Table 3). This *K*_*M*_ value in the low micromolar range indicates that the ABA3-STR18 interaction may be physiologically relevant.

### STR18 is persulfidated upon reaction with ABA3

As L-cysteine is not a sulfur donor for STR18 (Fig. S3), we assumed that STR18 stimulated ABA3 activity by reducing the persulfide formed on ABA3 more efficiently than the reductants used in the activity assay. In other words, this implied the transfer of sulfur atoms from ABA3 to STR18. To test this assumption, we analyzed by mass spectrometry the molecular mass of STR18 before and after incubation with a catalytic amount of ABA3 and an excess of L-cysteine. An increase of the molecular mass of STR18 by 31.3 Da, corresponding to the mass of a sulfur atom, was observed after the reaction as compared with a pre-reduced STR18. As this mass difference disappeared after a DTT treatment, we concluded that STR18 was mono-persulfidated upon reaction with ABA3 in the presence of L-cysteine (Table 4, Fig. S4).

**Table 4.** Electrospray ionization mass spectrometry analysis of the redox state of STR18 and its monocysteinic variants.

| **Protein** | **Theoretical mass (Da)** | **Theoretical mass without Met (Da)** | **Pre-reduced with DTT** | **Treatment with ABA3 and L-cysteine** | **$\Delta$ mass (Da)** |
| --- | --- | --- | --- | --- | --- |
| STR18 | 17,117.0 | 16,985.8 | 16,985.4 | 17,016.7 | 31.3 |
| STR18 C47S | 17,100.9 | 16,969.8 | 16,968.2 | 17,001.2 | 33.0 |
| STR18 C89S | 17,100.9 | 16,969.8 | 16,970.0 | 16,969.0 | -1.0 |

Reduced proteins and proteins incubated with L-cysteine and ABA3 were analyzed by mass spectrometry. The mass accuracy is generally $\pm 0.5$ -1 Da. Note that the mass decrease of *ca* 131 Da compared to the theoretical molecular masses indicated that the methionine was cleaved off in *E. coli*.

To firmly establish which cysteine of STR18 is persulfidated by ABA3, similar incubation of STR18 variants with ABA3 and L-cysteine have been performed and analyzed by mass spectrometry. A DTT-reversible increase of 33 Da was detected for the C47S variant but not the C89S variant (Table 4, Fig. S5, S6). This indicated that STR18 was persulfidated on Cys89. Altogether, these data demonstrated the persulfidation of the Cys89 of STR18 by ABA3 in the presence of L-cysteine and the dispensable role of Cys47 for both the TST activity and the ABA3-mediated persulfidation of STR18.

### STR18 promotes trans-persulfidation reaction between two proteins

In the absence of a known sulfur acceptor(s) for STR18, we have investigated the capability of STR18 to transfer a sulfur atom to a protein by using roGFP2. We have thus tested the oxidation of a pre-reduced roGFP2 in the presence of STR18 and thiosulfate (Fig. 4A). Whereas thiosulfate alone had no effect, the presence of STR18 promoted roGFP2 oxidation (Fig. 4A). This result validated a trans-persulfidation reaction between thiosulfate, STR18 and roGFP2. Then, we investigated roGFP2 oxidation by STR18 in the presence of ABA3 and L-cysteine (Fig. 4B). We first analyzed whether STR18 or ABA3 alone were able to oxidize roGFP2 with L-cysteine. The obtained results confirmed that L-cysteine is not a sulfur donor for STR18 and indicated that ABA3 is unable to promote roGFP2 oxidation. On the contrary, roGFP2 was oxidized by the whole sulfur relay system (L-cysteine, ABA3, STR18). Similar results were obtained using the STR18 C47S variant while the STR18 C89S did not promote the ABA3-dependent roGFP2 oxidation (Fig. 4B) These results demonstrated that STR18 mediates sulfur transfer from ABA3 to roGFP2, thus catalyzing a trans-persulfidation reaction between both proteins. As already observed for TST activity, only Cys89 is mandatory for the trans-persulfidation reaction catalyzed by STR18.

**Figure 4.**
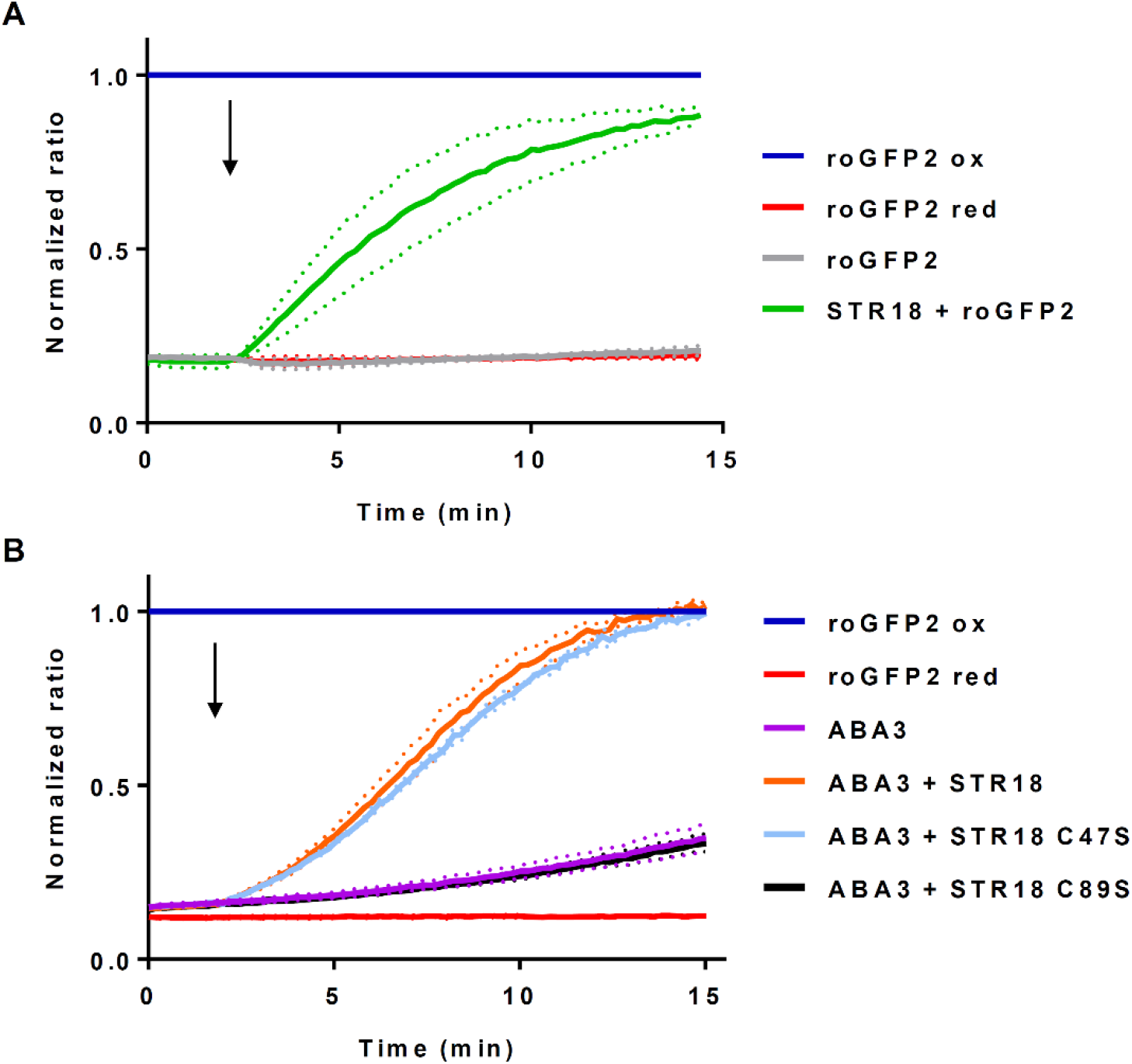
STR18 catalyzes the oxidation of roGFP2 via trans-persulfidation. A. Persulfide-dependent oxidation kinetics of 1 μM roGFP2 in the presence of 5 μM STR18 and 5 mM thiosulfate. B. Persulfide-dependent oxidation kinetics of 1 μM roGFP2 in the presence of 1mM L-cysteine, 1 mM PLP, 1 μM ABA3 and 5 μM STR18 or its cysteinic variants. The arrows indicate the addition of STR18, if present, after 2 min. The fully reduced or oxidized roGFP2 used as references were obtained after incubation with 10 mM DTT or H_2_O_2_ respectively. The ratio of 400/480 nm was normalized to the respective value of maximal roGFP2 oxidation by H_2_O_2_. The data are represented as mean ± SD of three independent experiments.

### A. thaliana STR18 and ABA3 interact in planta

To test whether an ABA3-STR18 interaction could be detected *in planta*, we performed split-luciferase complementation assays in transiently transformed tobacco leaves (Fig. 5). The bioluminescence emission corresponding to the activity of reconstituted luciferase was tested for different combinations with either candidates fused to the N-terminus of the N-terminal (nLuc) or to the C-terminus of C-terminal (cLuc) domain of luciferase. An intense luciferase signal was detected when both ABA3 and STR18 were fused either to nLuc or cLuc domains and co-expressed in tobacco leaves (Fig. 5). On the contrary, no signal was detected when ABA3 or STR18 were co-expressed with a subunit of the ATP citrate lyase (ACL1) known to be localized in cytosol like ABA3 and STR18 (Fig. 5). In combination with free-nLuc and free-cLuc controls, the latter finding confirmed the specificity of the bioluminescence signal detected after co-expression of ABA3 and STR18. Taken together, these results strongly suggest that ABA3 and STR18 interact in the cytosol of plant cells.

**Figure 5.**
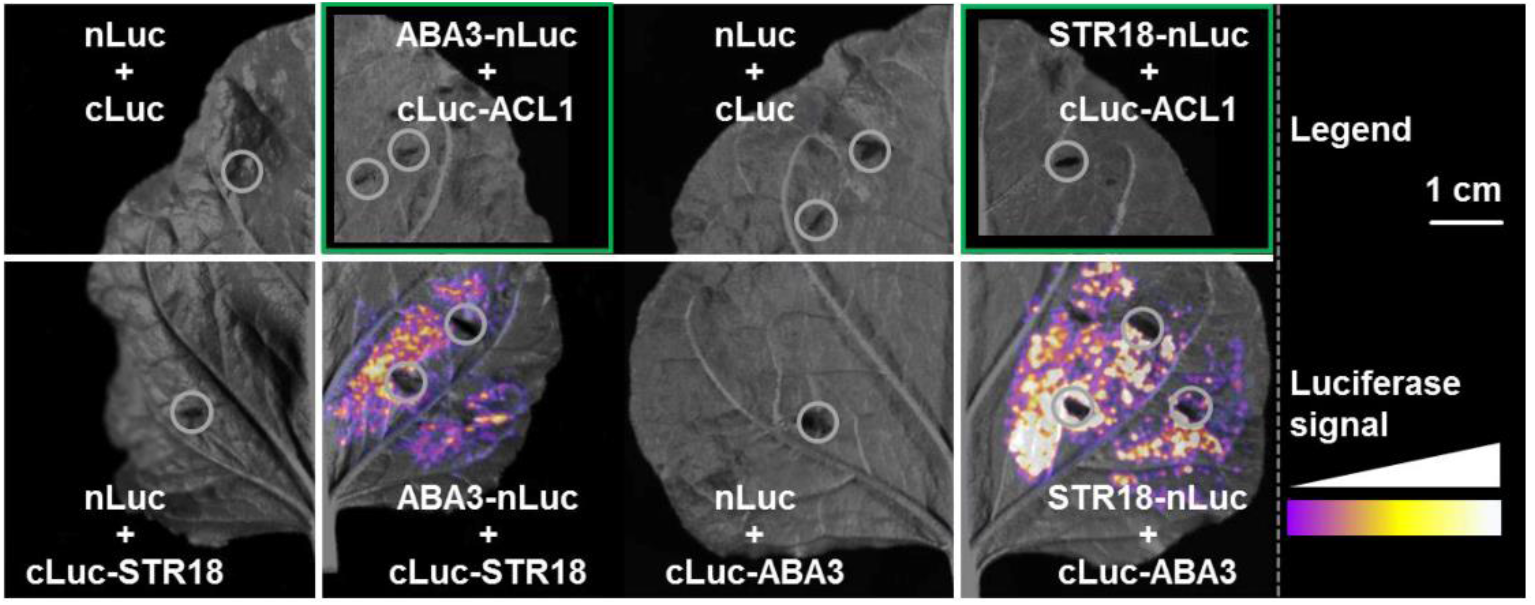
Split-luciferase complementation assays demonstrate close proximity of STR18 and ABA3 *in planta*. STR18, ABA3 and the negative control protein ACL1 were fused with the N-terminal (nLuc) or the C-terminal domain of luciferase (cLuc) to test for reassembling of enzymatically active luciferase based on the interaction of the respective fusion partners. DNA constructs encoding for the STR18 or ABA3 proteins in reciprocal fusion with both Luc domains were transiently transformed in *Nicotiana benthamiana* leaves, and tested for reconstitution of luciferase activity. Co-expression of nLuc, cLuc or the fusion of cLuc to ACL1 (cLuc-ACL1) served as negative controls. The detected signal intensity is shown in false color. Grey circles indicate inoculation sites of *Agrobacterium* for transient transformation. Green frames indicate digitally extracted pictures of individual leaves, allowing direct comparison of detected signals from different construct combinations. Scale bar: 1 cm.

## DISCUSSION

In plants, CDs are key enzymes involved in the maturation of both Fe-S- and Moco-containing proteins (14, 15, 19). As CDs act in the early steps of the maturation process and these metalloproteins fulfill important roles, the deletion of CD encoding genes in plants, more particularly *NFS1* and *NFS2*, is generally lethal or strongly affects development (14, 15). Considering that CDs serve as a central hub for sulfur mobilization and subsequent transfer to various metabolic pathways in non-photosynthetic organisms, we postulate that the strong phenotypes of mutant plants have thus so far prevented the identification of other sulfur-dependent pathways in which CDs are involved.

### The natural CD-Rhd fusion protein of Pseudorhodoferax sp. represents a highly efficient cysteine desulfurase isoform

The multiple properties of CD proteins are also evident from the existence of fusion proteins containing a CD domain associated with diverse protein domains. The plant ABA3 possesses a MOSC domain which links the protein function with Moco maturation. According to the known interaction between *E. coli* IscS and the ThiI or YnjE sulfurtransferases (26, 27), CD-Rhd chimera exist in several bacteria. Here, we described that a *Pseudorhodoferax* CD-Rhd is a PLP containing-homodimer exhibiting a dual activity profile, as it catalyzes cysteine and thiosulfate desulfuration using its respective functional domains. Noteworthy, this *Pseudorhodoferax* CD-Rhd is the most efficient cysteine desulfurase characterized so far, with a rate of sulfide formation of 7600 nmol min^-1^ mg^-1^ in the presence of L-cysteine and DTT. This activity is between 20 and 600-fold higher as compared with bacterial (*A. vinelandii* NifS and IscS, *B. subtilis* SufS, *E. coli* IscS and SufS, *Erwinia chrysanthemi* SufS), and eukaryotic CDs *(A. thaliana* NFS2, human and yeast NFS1) (Table 5). This remains true if we consider the activity of group II CD members in the presence of their respective activators. Indeed, the rate of sulfide formation ranged from 550 nmol min^-1^ mg^-1^ for *A. thaliana* NFS2-SUFE1, to 750 and 900 nmol min^-1^ mg^-1^ for *E. chrysanthemi* and *E. coli* SufS-SufE, respectively (Table 5) (11, 16). The CD activity of the CD-Rhd C466S variant, for which the Rhd domain is inactive, remains high despite it decreased by ∼10-fold in the presence of GSH and β-ME compared to CD-Rhd. In this case, with a rate of sulfide formation of 553 nmol min^-1^ mg^-1^, this CD activity is in the same range as those determined for *E. coli* IscS, SufS-SufE and *A. thaliana* NFS2-SUFE1 (Table 5). All these results indicate that (i) the CD domain of *Pseudorhodoferax* CD-Rhd is very active and (ii) the fusion to a Rhd domain increases its activity with the catalytic cysteine of the Rhd domain acting as a sulfur acceptor, as observed for group II CD isoforms and their specific activators.

**Table 5.**
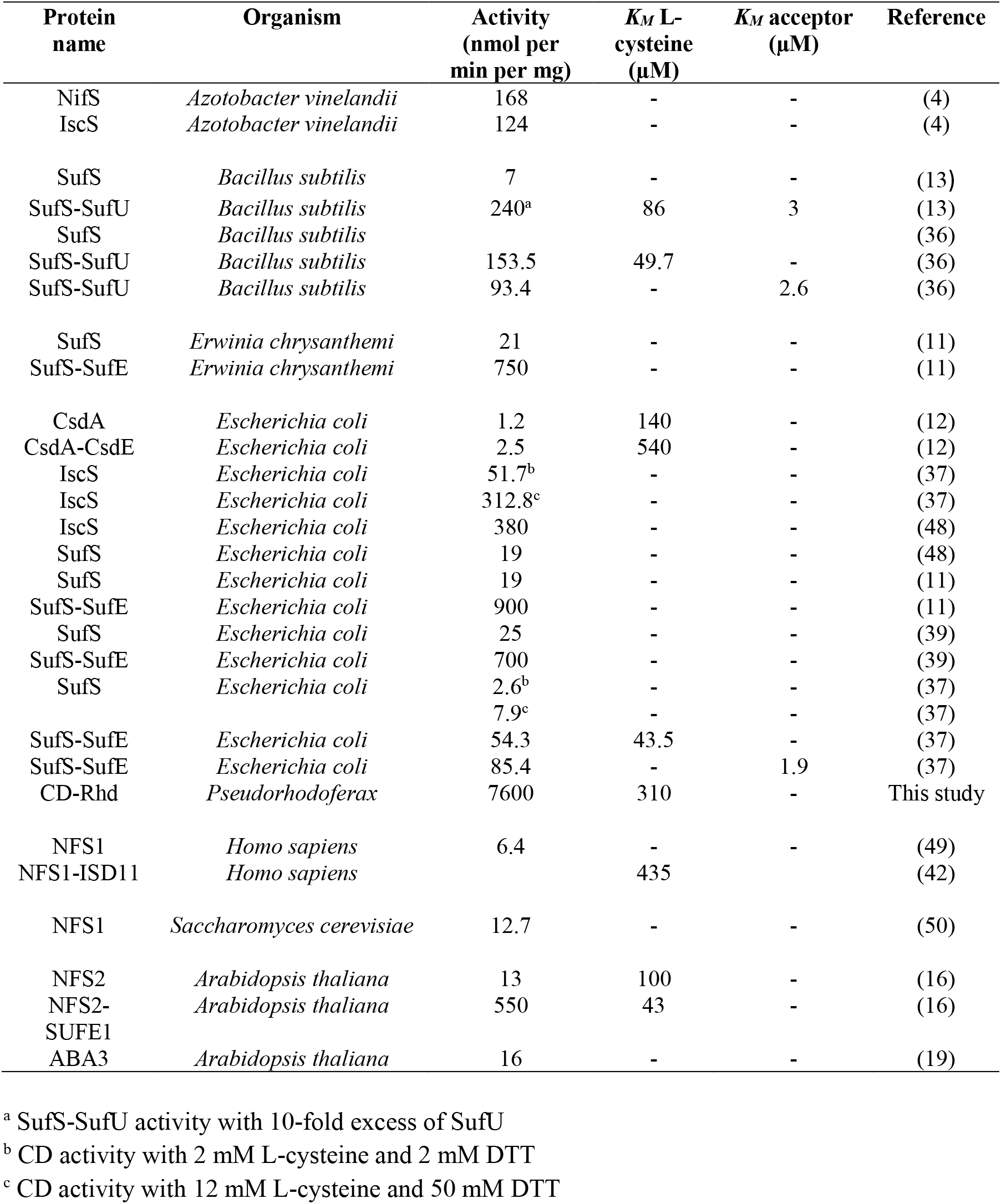
Catalytic properties of characterized CD isoforms from various organisms.

*Pseudorhodoferax* CD-Rhd exhibits a TST activity indicating that the Rhd domain is also functional. It displayed a better affinity for thiosulfate as compared with *E. coli* TST isoforms, GlpE and PspE, (*K*_*M*_,_*app*_ of 756 μM *vs* 78 mM and 2.7 mM) (34, 35) but a 4-fold lower catalytic efficiency than STR18 (Table 2). Hence, CD-Rhd is a bifunctional enzyme using both L-cysteine and thiosulfate as sulfur donors. Nevertheless, considering catalytic efficiencies of both CD and TST activities (2.2 × 10^4^ M^-1^ s^-1^ *vs* 2.7 × 10^3^ M^-1^ s^-1^ in the presence of β-ME) and the fast and specific CD domain-dependent oxidation of roGFP2, L-cysteine and the associated CD activity represent the preferential substrate and activity of *Pseudorhodoferax* CD-Rhd.

The efficient cysteine-dependent oxidation of roGFP2 through trans-persulfidation reaction catalyzed by CD-Rhd (Fig. 2) also suggests that a role in persulfidation of target proteins may be physiologically relevant. Moreover, considering H_2_S release measured in the presence of various reductants and notably GSH, *Pseudorhodoferax* CD-Rhd might be also involved in the synthesis of H_2_S and/or of low-molecular-weight persulfides.

### ABA3-STR18 represents a new cytosolic pathway of sulfur trafficking in plant cells

The existence of such natural fusion proteins prompted us to analyze whether the CD activity of ABA3 is enhanced by a STR or in other words if a persulfide transfer reaction is possible between these proteins. In the presence of L-cysteine and DTT, ABA3 displayed an activity and *k*_*cat*_ value in the range of the values reported for other CD isoforms (Table 5) (21). Concerning the impact of STR18, the catalytic efficiency of ABA3, measured under steady-state conditions, increased 5-fold in the presence of STR18, an effect mostly due to a 3-fold lower apparent *K*_*M*_ value for L-cysteine. Similar effects were reported for the plastidial SUFE1 protein which decreased by a factor 2 the *K*_*M*_ value of NFS2 for L-cysteine and increased 42-fold the rate of sulfide formation by NFS2 (16). Furthermore, the low *K*_*M*_ value of 1.2 μM of ABA3 for STR18 determined under steady-state conditions is consistent with the values obtained for the *B. subtilis* SufS-SufU and *E. coli* SufS-SufE couples (Table 5) (13, 36, 37). Interestingly, in all these examples the apparent *K*_*M*_ values of the CDs for their protein partners are lower than their apparent *K*_*M*_ values for L-cysteine (8-fold lower for ABA3-STR18 and 20-fold lower for *B. subtilis* SufS-SufU and *E. coli* SufS-SufE couples) (13, 36, 37). The physical interaction between both proteins observed with split-luciferase complementation suggests a specific, physiologically relevant ABA3-STR18 interaction.

### ABA3-STR18 couple catalyzes trans-persulfidation reactions

Both the TST activity and the positive effect on ABA3 activity of STR18 underlined the ability of STR18 to form an intermediate persulfide as demonstrated previously for *A. vinelandii* Rhd isoform RhdA in the presence of *E. coli* IscS (38). This was also expected from the sulfur transfer observed from *E. coli* SufS and CsdA to SufE and CsdE respectively (10, 39). The ABA3-dependent persulfidation of the catalytic Cys89 of STR18 was indeed demonstrated by mass spectrometry after incubation of pre-reduced STR18 with both L-cysteine and ABA3 (Table 4). By accepting the sulfur atom, STR18 stimulates the CD activity of ABA3 and regenerates its active form being able to bind the next cysteine molecule (40). In the absence of known STR18 partner(s), we further demonstrated the capacity of STR18 to perform trans-persulfidation reactions from either thiosulfate or ABA3 and L-cysteine to roGFP2 (Fig. 4). From an experimental point of view, the roGFP2 assay enables us to study the ability of a candidate protein to catalyze trans-persulfidation reaction in the absence of known partners. It was recently demonstrated that STR1 and STR2, which possess two Rhd domains, efficiently transfer a persulfide to roGFP2 (30). Arabidopsis STR16, another single Rhd-domain containing protein, is also able to catalyze roGFP2 oxidation in the presence of thiosulfate (Fig. S7). All these results suggest that the catalysis of trans-persulfidation reaction might be a conserved function of STRs. From the apparent *K*_*M*_ values of STR18 for thiosulfate (527 ± 8 μM) and of ABA3 for STR18 (1.2 ± 0.2 μM), and the ability of ABA3 to promote STR18 persulfidation more efficiently than thiosulfate, ABA3 may be seen as the preferential sulfur donor for STR18.

### Relationships between ABA3-STR18 and other cytosolic STR isoforms

All these results represent the first evidence of a functional relationship between CD and STR in plant cells. Of interest, the sulfur transfer pathway from ABA3 to STR18 may be independent of a sulfur transfer to the MOCS domain and thus independent of Moco sulfuration. In which physiological context such a pathway is relevant remains to be demonstrated because other cytosolic STRs are present in Arabidopsis. In addition to STR18, *A. thaliana* possesses at least two other cytosolic STR isoforms, the MST isoform STR2 and the two domain-containing protein STR13 also referred to as CNX5/MOCS3 (24). Noteworthy, STR2 and STR13 are present in all eukaryotic photosynthetic organisms whereas STR18 is present only in dicotyledonous plants (24). The physiological function(s) of STR2 and STR18 are yet unknown *in planta* unlike STR13, which possesses a dual function, delivering the sulfur needed for the thio-modification of cytosolic tRNAs and for Moco biosynthesis owing to its N-terminal domain (31, 41). In human cells, a cytosolic form of NFS1 was proposed to provide sulfur to MOCS3 eventually involving a relay by TUM1, the ortholog of plant STR2 (28, 42, 43).

While similar actors are present in plants, it may be that this cytosolic sulfur trafficking pathway is different between human and plant cells. Indeed, *A. thaliana str2* null mutant lines have no phenotype whereas *str13* null mutants (*cnx5-1* and *cnx5-2*) are sterile and exhibit a severe dwarf phenotype with slightly green and morphologically aberrant leaves (41, 44). On the contrary, *aba3* mutants (*aba3-1, aba3-2, los5-1, los5-2, aba3-7, aba3-8*) have distinct and less severe phenotypes (45–47) than *str13* mutants. This suggests either that STR13 persulfidation would be independent of ABA3 or that STR13 possesses additional functions.

From these results, we propose that in addition to its role in the maturation of the Moco-containing proteins, XDH and AO, ABA3 acts as a sulfur donor to STR proteins (either STR18 as demonstrated here or other cytosolic members such as STR2). The trans-persulfidation pathway involving cysteine and an ABA3-STR couple might thus represent an uncharacterized sulfur trafficking pathway in the cytosol of plants.

## EXPERIMENTAL PROCEDURES

### Materials

3-MP (sodium salt) was purchased from Santa Cruz Biotechnology (Dallas, TX, USA), lead (II) acetate, L-cysteine, thiosulfate, GSH and β-mercaptoethanol were from Sigma-Aldrich (St Louis, MO, USA).

### Cloning and site-directed mutagenesis

The sequences coding for *A. thaliana* STR16 (At5g66040) STR18 (At5g66170), the full-length ABA3 (At1g16540) were cloned into the *Nde*I and *Bam*HI restriction sites of pET15b. Catalytic cysteine (Cys80) of STR16 and both cysteine residues (Cys47 and Cys89) of STR18 were individually substituted into serines to generate pET15b-STR16 C80S, pET15b-STR18 C47S and pET15b-STR18 C89S recombinant plasmids. A synthetic cDNA (Genecust, France) coding for CD-Rhd fusion protein (WP_056898193.1) from *Pseudorhodoferax sp*. Leaf274 was cloned into the *Nde*I and *Bam*HI restriction sites of pET15b. The cysteine in position 466 was substituted to serine to generate a pET15b-CD-Rhd C466S recombinant plasmid. All primers used in this study are listed in Table S1.

### Heterologous expression in E. coli and purification of recombinant proteins

For protein expression, the *E. coli* BL21 (DE3), C41 (DE3) and Rosetta2 (DE3) pLysS strains were transformed respectively with pET15b At*STR16*, At*STR18*, At*ABA3, Pseudorhodoferax* CD-Rhd and CD-Rhd C466S. The BL21 (DE3) and C41 (DE3) strains also contained the pSBET plasmid which allows expression of the tRNA needed to recognize the AGG and AGA rare codons. Cell cultures were progressively amplified up to 2.4 l, for STR16, STR16 C80S, STR18, STR18 C47S, STR18 C89S, CD-Rhd and CD-Rhd C466S, and 4.8 l for ABA3, in LB medium supplemented with 50 μg/ml of ampicillin and kanamycin for BL21 and C41 strains or with 50 μg/ml of ampicillin and 34 μg/ml of chloramphenicol for Rosetta2 strain and grown at 37°C. STR18 expression was induced at exponential phase by adding 100 μM isopropyl β-D-thiogalactopyranoside (IPTG) for 4 h at 37°C. For ABA3, CD-Rhd and CD-Rhd C466S, the culture protocol was modified. At exponential phase, the cultures were supplemented with ethanol 0.5% (v:v) and 100 μM pyridoxine hydrochloride and placed at 4°C for 2 h. Protein expression was then induced by adding 100 μM IPTG for 18 h at 20°C. After centrifugation (20 min at 6,380 x g), the cell pellets were resuspended in about 20 ml of 50 mM Tris-HCl pH 8.0, 300 mM NaCl, 10 mM imidazole buffer and stored at -20°C.

Cell lysis was completed by sonication (3 × 1 min with intervals of 1 min) and the soluble and insoluble fractions were separated by centrifugation for 30 min at 27,216 x g. For all proteins, the soluble fraction was loaded on Ni^2+^ affinity column (Sigma-Aldrich, St Louis MO, USA). After extensive washing, proteins were eluted by a 50 mM Tris-HCl pH 8.0, 300 mM NaCl, 250 mM imidazole buffer. The recombinant proteins were concentrated by ultrafiltration under nitrogen pressure and dialyzed (Amicon, YM10 membrane), and finally stored in a 30 mM Tris-HCl pH 8.0, 200 mM NaCl buffer supplemented with 5 mM DTT and 50% glycerol at -20°C. Protein concentrations were determined spectrophotometrically using a molecular extinction coefficient at 280 nm of 10,095 M^-1^ cm^-1^ for STR16 and 9970 M^-1^ cm^-1^ for its monocysteinic variant, 11,585 M^-1^ cm^-1^ for STR18 and 11,460 M^-1^ cm^-1^ for its monocysteinic variants, 97,845 M^-1^ cm^-1^ and 57,800 M^-1^ cm^-1^, for ABA3, and 47,690 M^-1^ cm^-1^ for CD-Rhd and CD-Rhd C466S, respectively. The roGFP2 recombinant protein used in this study has been purified as described previously (51).

### Determination of the oligomerization state of CD-Rhd

The oligomerization state of CD-Rhd and CD-Rhd C466S variant was analyzed by analytical size-exclusion chromatography as described previously (52). The detection was recorded by measuring absorbances at 280 and 418 nm. The column was calibrated using the following molecular weight standards: thyroglobulin (669 kDa, 8.8 ml), β-amylase (200 kDa, 12 ml), bovine serum albumin (66 kDa, 13.6 ml) and cytochrome c (12.4 kDa, 16.8 ml).

### Cysteine desulfurase activity assays

The CD activity was assayed at 25°C in a final volume of 400 μl of 30 mM Tris-HCl pH 8.0 buffer, 10 μM PLP, 5 mM reductant (DTT, GSH or β-mercaptoethanol) and 10 nM CD-Rhd, 100 nM CD-Rhd C466S or 1 μM ABA3. To assess the impact of STR18 on ABA3 activity, 5 μM STR18 was added in the reaction mixture. The reaction was initiated by adding L-cysteine and stopped after 30 min by adding 50 μl of 20 mM N,N-dimethyl-p-phenylenediamine dihydrochloride (DMPD, prepared in 7.2 M HCl). The addition of 50 μl of 30 mM FeCl_3_ (prepared in 1.2 M HCl), followed by a 20 min incubation led to formation of methylene blue, which was then measured at 670 nm. Sodium sulfide (Na_2_S) in the range of 1 to 100 μM was used for standard curve calibration.

### Thiosulfate sulfurtransferase activity assays

The thiosulfate sulfurtransferase activity of CD-Rhd, STR18 and their variants was assayed at 25°C in a final volume of 500 μl of 30 mM Tris-HCl pH 8.0 buffer, 5 mM β-mercaptoethanol, 0.4 mM lead (II) acetate (Sigma-Aldrich, St Louis MO, USA), various concentrations of thiosulfate ranging from 0 to 5 mM and 100 nM enzyme. The reaction was initiated by adding CD-Rhd or STR18 and the rate of lead sulfide formation was monitored at 390 nm using a molar extinction coefficient of 5,500 M^−1^ cm^−1^.

### Detection of persulfidated STR18 by mass spectrometry

In a final volume of 150 μl of 30 mM Tris HCl pH 8.0, 200 mM NaCl buffer, 150 μM of pre-reduced STR18, STR18 C47S and STR18 C89S were incubated 30 min in the presence of 300 μM L-cysteine, 2 μM ABA3 and 5 μM PLP at 25°C. After extensive dialysis, samples were split in two parts and treated or not with 1 mM DTT. Mass spectrometry analysis of these samples was performed using a Bruker microTOF-Q spectrometer (Bruker Daltonik, Bremen, Germany), equipped with Apollo II electrospray ionization source with ion funnel, operated in the negative ion mode. The concentrated samples in formic acid were injected at a flow rate of 10-20 μl min^-1^. The potential between the spray needle and the orifice was set to 4.5 kV. Before each run the instrument was calibrated externally with the Tunemix™ mixture (Agilent Technologies, Santa Clara, CA, USA) in quadratic regression mode. Data were analyzed with the DataAnalysis software (Bruker).

### roGFP2 oxidation experiments

The capacity of CD-Rhd, ABA3, STR16 and STR18 to oxidize roGFP2 was analyzed *in vitro* by ratiometric time-course measurements on a fluorescence plate reader (EnSight multimode plate reader, PerkinElmer) with excitation at 400 ± 10 and 480 ± 10 nm and detection of emitted light at 520 nm with a bandwidth of 10 nm. The maximum oxidation and reduction of roGFP2 were defined using 10 mM H_2_O_2_ and DTT. Pre-reduced roGFP2 was obtained by incubation with 10 mM DTT for 1 h and subsequent desalting on a G25 column to remove excess DTT. In a final volume of 400 μl of 30 mM Tris-HCl pH 8.0, 200 mM NaCl, the reaction mixtures contained 1 μM pre-reduced roGFP2 and either 5 mM thiosulfate and 5 μM STR16/STR18/CD-Rhd or 1 mM L-cysteine, 5 μM CD-Rhd or 1 mM L-cysteine, 10 μM PLP, 1 μM ABA3 and 5 μM STR18.

### Split-luciferase complementation assays

Full-length coding sequences of ABA3, STR18 and the negative control ACL1 (At1g10670) were selectively amplified with primers defined in Table S1 and cloned via *Kpn*I and *Sal*I restriction endonucleases into the pCAMBIA1300-cLuc or the pCAMBIA1300-nLuc vectors described in (53). The resulting fusion constructs were named cLuc-ABA3, cLuc-STR18, ABA3-nLuc, STR18-nLuc and cLuc-ACL1. Different combinations of cLuc-fusion and nLuc-fusion constructs were co-expressed in tobacco (*Nicotiana benthamiana*) leaves after *Agrobacterium*-mediated transient transformation according to (54). After *Agrobacterium* inoculation, plants were kept for 24 hours in the dark and subsequently grown for two days under long day conditions (16 h light 250 μE, 8 h dark, temperature 25°C, humidity: 50%) to allow expression of the protein of interest in fusion with the N-terminal (nLuc) or C-terminal (cLuc) fragment of luciferase. The abaxial sides of the transformed leaves were sprayed with luciferase substrate (1 mM luciferin) and the substrate was allowed to enter the leaf for 5 min. The resulting luciferase signal was detected with the digital camera system ‘ImageQuant LAS 4000’ (GE Healthcare, Chicago, IL, USA) and visualized with the open access software suite ‘Image J’ (National Institutes of Health, Bethesda, MD, USA).

## Data availability

All data are presented in the manuscript.

## Supporting information

This article contains supporting information.

## Acknowledgments

Technical support from Fabien Lachaud and François Dupire of the “Service Commun de Spectrométrie de Masse et Chromatographie” of the Université de Lorraine is gratefully acknowledged.

## Author contributions

B.S. and J.C. conceived the research with specific input from M.W. and N.R. B.S., A.M., D.C., S.K.S., T.D., M.Z. and M.H. performed the experiments. M.W., N.R. and J.C. supervised the experiments and interpreted the data. B.S. and J.C. wrote the manuscript with support from A.M., D.C., M.W. and N.R. All authors read and approved the manuscript.

## Funding and additional information

This work was supported by the Agence Nationale de la Recherche as part of the “Investissements d’Avenir” program (ANR-11-LABX-0002-01, Lab of Excellence ARBRE), by grant no. ANR-16-CE20-0012 and by ECOS-*Sud* program. A.M. is recipient of a Feodor Lynen Research Fellowship and S.K.S. is a recipient of Humboldt Research Fellowship from the Alexander von Humboldt Foundation. The PhD salary of D.C. was provided by a funding from the Lorraine University of Excellence (LUE).

## REFERENCES

1. Mueller, E. G. (2006) Trafficking in persulfides: delivering sulfur in biosynthetic pathways. Nat. Chem. Biol. 2, 185–194

2. Behshad, E., and Bollinger, J. M. (2009) Kinetic analysis of cysteine desulfurase CD0387 from Synechocystis sp. PCC 6803: formation of the persulfide intermediate. Biochemistry. 48, 12014–12023

3. Zheng, L., White, R. H., Cash, V. L., Jack, R. F., and Dean, D. R. (1993) Cysteine desulfurase activity indicates a role for NIFS in metallocluster biosynthesis. Proc. Natl. Acad. Sci. U. S. A. 90, 2754–2758

4. Zheng, L., Cash, V. L., Flint, D. H., and Dean, D. R. (1998) Assembly of iron-sulfur clusters identification of an iscSUA-hscBA-fdx gene cluster from Azotobacter vinelandii. J. Biol. Chem. 273, 13264–13272

5. Patzer, S. I., and Hantke, K. (1999) SufS is a NifS-like protein, and SufD is necessary for stability of the [2Fe-2S] FhuF protein in Escherichia coli. J. Bacteriol. 181, 3307–3309

6. Mihara, H., and Esaki, N. (2002) Bacterial cysteine desulfurases: their function and mechanisms. Appl. Microbiol. Biotechnol. 60, 12–23

7. Shi, R., Proteau, A., Villarroya, M., Moukadiri, I., Zhang, L., Trempe, J.-F., Matte, A., Armengod, M. E., and Cygler, M. (2010) Structural basis for Fe–S cluster assembly and tRNA thiolation mediated by IscS protein–protein interactions. PLoS Biol. 8, e1000354

8. Roret, T., Pégeot, H., Couturier, J., Mulliert, G., Rouhier, N., and Didierjean, C. (2014) X-ray structures of Nfs2, the plastidial cysteine desulfurase from Arabidopsis thaliana. Acta Crystallogr. Sect. F Struct. Biol. Commun. 70, 1180–1185

9. Black, K. A., and Dos Santos, P. C. (2015) Shared-intermediates in the biosynthesis of thio-cofactors: Mechanism and functions of cysteine desulfurases and sulfur acceptors. Biochim. Biophys. Acta. 1853, 1470–1480

10. Outten, F. W., Wood, M. J., Munoz, F. M., and Storz, G. (2003) The SufE protein and the SufBCD complex enhance SufS cysteine desulfurase activity as part of a sulfur transfer pathway for Fe-S cluster assembly in Escherichia coli. J. Biol. Chem. 278, 45713–45719

11. Loiseau, L., Ollagnier-de-Choudens, S., Nachin, L., Fontecave, M., and Barras, F. (2003) Biogenesis of Fe-S cluster by the bacterial Suf system: SufS and SufE form a new type of cysteine desulfurase. J. Biol. Chem. 278, 38352–38359

12. Loiseau, L., Ollagnier-de Choudens, S., Lascoux, D., Forest, E., Fontecave, M., and Barras, F. (2005) Analysis of the heteromeric CsdA-CsdE cysteine desulfurase, assisting Fe-S cluster biogenesis in Escherichia coli. J. Biol. Chem. 280, 26760–26769

13. Selbach, B., Earles, E., and Dos Santos, P. C. (2010) Kinetic analysis of the bisubstrate cysteine desulfurase SufS from Bacillus subtilis. Biochemistry. 49, 8794–8802

14. Frazzon, A. P. G., Ramirez, M. V., Warek, U., Balk, J., Frazzon, J., Dean, D. R., and Winkel, B. S. J. (2007) Functional analysis of Arabidopsis genes involved in mitochondrial iron-sulfur cluster assembly. Plant Mol. Biol. 64, 225–240

15. Van Hoewyk, D., Abdel-Ghany, S. E., Cohu, C. M., Herbert, S. K., Kugrens, P., Pilon, M., and Pilon-Smits, E. A. H. (2007) Chloroplast iron-sulfur cluster protein maturation requires the essential cysteine desulfurase CpNifS. Proc. Natl. Acad. Sci. U. S. A. 104, 5686–5691

16. Ye, H., Abdel-Ghany, S. E., Anderson, T. D., Pilon-Smits, E. A. H., and Pilon, M. (2006) CpSufE activates the cysteine desulfurase CpNifS for chloroplastic Fe-S cluster formation. J. Biol. Chem. 281, 8958–8969

17. Murthy, N. M. U., Ollagnier-de-Choudens, S., Sanakis, Y., Abdel-Ghany, S. E., Rousset, C., Ye, H., Fontecave, M., Pilon-Smits, E. A. H., and Pilon, M. (2007) Characterization of Arabidopsis thaliana SufE2 and SufE3: functions in chloroplast iron-sulfur cluster assembly and Nad synthesis. J. Biol. Chem. 282, 18254–18264

18. Xu, X. M., and Møller, S. G. (2006) AtSufE is an essential activator of plastidic and mitochondrial desulfurases in Arabidopsis. EMBO J. 25, 900–909

19. Bittner, F., Oreb, M., and Mendel, R. R. (2001) ABA3 is a molybdenum cofactor sulfurase required for activation of aldehyde oxidase and xanthine dehydrogenase in Arabidopsis thaliana. J. Biol. Chem. 276, 40381–40384

20. Kaufholdt, D., Baillie, C.-K., Meyer, M. H., Schwich, O. D., Timmerer, U. L., Tobias, L., van Thiel, D., Hänsch, R., and Mendel, R. R. (2016) Identification of a protein-protein interaction network downstream of molybdenum cofactor biosynthesis in Arabidopsis thaliana. J. Plant Physiol. 207, 42–50

21. Heidenreich, T., Wollers, S., Mendel, R. R., and Bittner, F. (2005) Characterization of the NifS-like domain of ABA3 from Arabidopsis thaliana provides insight into the mechanism of molybdenum cofactor sulfuration. J. Biol. Chem. 280, 4213–4218

22. Wollers, S., Heidenreich, T., Zarepour, M., Zachmann, D., Kraft, C., Zhao, Y., Mendel, R. R., and Bittner, F. (2008) Binding of sulfurated molybdenum cofactor to the C-terminal domain of ABA3 from Arabidopsis thaliana provides insight into the mechanism of molybdenum cofactor sulfuration. J. Biol. Chem. 283, 9642–9650

23. Bordo, D., and Bork, P. (2002) The rhodanese/Cdc25 phosphatase superfamily. Sequence-structure-function relations. EMBO Rep. 3, 741–746

24. Moseler, A., Selles, B., Rouhier, N., and Couturier, J. (2020) Novel insights into the diversity of the sulfurtransferase family in photosynthetic organisms with emphasis on oak. New Phytol. 226, 967–977

25. Cipollone, R., Ascenzi, P., and Visca, P. (2007) Common themes and variations in the rhodanese superfamily. IUBMB Life. 59, 51–59

26. Dahl, J.-U., Urban, A., Bolte, A., Sriyabhaya, P., Donahue, J. L., Nimtz, M., Larson, T. J., and Leimkühler, S. (2011) The identification of a novel protein involved in molybdenum cofactor biosynthesis in Escherichia coli. J. Biol. Chem. 286, 35801–35812

27. Kambampati, R., and Lauhon, C. T. (2000) Evidence for the transfer of sulfane sulfur from IscS to ThiI during the in vitro biosynthesis of 4-thiouridine in Escherichia coli tRNA. J. Biol. Chem. 275, 10727–10730

28. Fräsdorf, B., Radon, C., and Leimkühler, S. (2014) Characterization and interaction studies of two isoforms of the dual localized 3-mercaptopyruvate sulfurtransferase TUM1 from humans. J. Biol. Chem. 289, 34543–34556

29. Noma, A., Sakaguchi, Y., and Suzuki, T. (2009) Mechanistic characterization of the sulfur-relay system for eukaryotic 2-thiouridine biogenesis at tRNA wobble positions. Nucleic Acids Res. 37, 1335–1352

30. Moseler, A., Dhalleine, T., Rouhier, N., and Couturier, J. (2021) Arabidopsis thaliana 3-mercaptopyruvate sulfurtransferases interact with and are protected by reducing systems. J. Biol. Chem. 296, 100429

31. Bauer, M., Dietrich, C., Nowak, K., Sierralta, W. D., and Papenbrock, J. (2004) Intracellular localization of Arabidopsis sulfurtransferases. Plant Physiol. 135, 916–926

32. Selles, B., Moseler, A., Rouhier, N., and Couturier, J. (2019) Rhodanese domain-containing sulfurtransferases: multifaceted proteins involved in sulfur trafficking in plants. J. Exp. Bot. 70, 4139–4154

33. Henne, M., König, N., Triulzi, T., Baroni, S., Forlani, F., Scheibe, R., and Papenbrock, J. (2015) Sulfurtransferase and thioredoxin specifically interact as demonstrated by bimolecular fluorescence complementation analysis and biochemical tests. FEBS Open Bio. 5, 832–843

34. Cheng, H., Donahue, J.L., Battle, S.E., Ray, W.K., and Larson, T.J. (2008) Biochemical and genetic characterization of PspE and GlpE, two single-domain sulfurtransferases of Escherichia coli. Open Microbiol. J. 2, 18–28

35. Ray, W. K., Zeng, G., Potters, M. B., Mansuri, A. M., and Larson, T. J. (2000) Characterization of a 12-Kilodalton rhodanese encoded by glpE of Escherichia coli and its Interaction with thioredoxin. J. Bacteriol. 182, 2277–2284

36. Albrecht, A. G., Peuckert, F., Landmann, H., Miethke, M., Seubert, A., and Marahiel, M. A. (2011) Mechanistic characterization of sulfur transfer from cysteine desulfurase SufS to the iron-sulfur scaffold SufU in Bacillus subtilis. FEBS Lett. 585, 465–470

37. Dai, Y., and Outten, F. W. (2012) The E. coli SufS-SufE sulfur transfer system is more resistant to oxidative stress than IscS-IscU. FEBS Lett. 586, 4016–4022

38. Forlani, F., Cereda, A., Freuer, A., Nimtz, M., Leimkühler, S., and Pagani, S. (2005) The cysteine-desulfurase IscS promotes the production of the rhodanese RhdA in the persulfurated form. FEBS Lett. 579, 6786–6790

39. Ollagnier-de-Choudens, S., Lascoux, D., Loiseau, L., Barras, F., Forest, E., and Fontecave, M. (2003) Mechanistic studies of the SufS-SufE cysteine desulfurase: evidence for sulfur transfer from SufS to SufE. FEBS Lett. 555, 263–267

40. Lehrke, M., Rump, S., Heidenreich, T., Wissing, J., Mendel, R. R., and Bittner, F. (2012) Identification of persulfide-binding and disulfide-forming cysteine residues in the NifS-like domain of the molybdenum cofactor sulfurase ABA3 by cysteine-scanning mutagenesis. Biochem. J. 441, 823–839

41. Nakai, Y., Harada, A., Hashiguchi, Y., Nakai, M., and Hayashi, H. (2012) Arabidopsis molybdopterin biosynthesis protein Cnx5 collaborates with the ubiquitin-like protein Urm11 in the thio-modification of tRNA. J. Biol. Chem. 287, 30874–30884

42. Marelja, Z., Stöcklein, W., Nimtz, M., and Leimkühler, S. (2008) A novel role for human Nfs1 in the cytoplasm: Nfs1 acts as a sulfur donor for MOCS3, a protein involved in molybdenum cofactor biosynthesis. J. Biol. Chem. 283, 25178–25185

43. Marelja, Z., Mullick Chowdhury, M., Dosche, C., Hille, C., Baumann, O., Löhmannsröben, H.-G., and Leimkühler, S. (2013) The L-cysteineteine desulfurase NFS1 is localized in the cytosol where it provides the sulfur for molybdenum cofactor biosynthesis in humans. PloS One. 8, e60869

44. Mao, G., Wang, R., Guan, Y., Liu, Y., and Zhang, S. (2011) Sulfurtransferases 1 and 2 play essential roles in embryo and seed development in Arabidopsis thaliana. J. Biol. Chem. 286, 7548–7557

45. Xiong, L., Ishitani, M., Lee, H., and Zhu, J. K. (2001) The Arabidopsis LOS5/ABA3 locus encodes a molybdenum cofactor sulfurase and modulates cold stress-and osmotic stress-responsive gene expression. Plant Cell. 13, 2063–2083

46. Zhong, R., Thompson, J., Ottesen, E., and Lamppa, G. K. (2010) A forward genetic screen to explore chloroplast protein import in vivo identifies Moco sulfurase, pivotal for ABA and IAA biosynthesis and purine turnover. Plant J. 63, 44–59

47. Watanabe, S., Sato, M., Sawada, Y., Tanaka, M., Matsui, A., Kanno, Y., Hirai, M. Y., Seki, M., Sakamoto, A., and Seo, M. (2018) Arabidopsis molybdenum cofactor sulfurase ABA3 contributes to anthocyanin accumulation and oxidative stress tolerance in ABA-dependent and independent ways. Sci. Rep. 8, 16592

48. Mihara, H., Kurihara, T., Yoshimura, T., and Esaki, N. (2000) Kinetic and mutational studies of three NifS homologs from Escherichia coli: mechanistic difference between L-cysteineteine desulfurase and L-selenocysteine lyase reactions. J. Biochem. (Tokyo). 127, 559–567

49. Li, K., Tong, W.-H., Hughes, R. M., and Rouault, T. A. (2006) Roles of the Mammalian Cytosolic Cysteine Desulfurase, ISCS, and Scaffold Protein, ISCU, in Iron-Sulfur Cluster Assembly. J. Biol. Chem. 281, 12344–12351

50. Mühlenhoff, U., Balk, J., Richhardt, N., Kaiser, J. T., Sipos, K., Kispal, G., and Lill, R. (2004) Functional characterization of the eukaryotic cysteine desulfurase Nfs1p from Saccharomyces cerevisiae. J. Biol. Chem. 279, 36906–36915

51. Meyer, A. J., Brach, T., Marty, L., Kreye, S., Rouhier, N., Jacquot, J.-P., and Hell, R. (2007) Redox-sensitive GFP in Arabidopsis thaliana is a quantitative biosensor for the redox potential of the cellular glutathione redox buffer. Plant J. 52, 973–986

52. Zannini, F., Roret, T., Przybyla-Toscano, J., Dhalleine, T., Rouhier, N., and Couturier, J. (2018) Mitochondrial Arabidopsis thaliana TRXo isoforms bind an iron−sulfur cluster and reduce NFU proteins in vitro. Antioxidants 7, 142

53. Chen, H., Zou, Y., Shang, Y., Lin, H., Wang, Y., Cai, R., Tang., X., and Zhou, J. M. (2008) Firefly luciferase complementation imaging assay for protein-protein interactions in plants. Plant Physiol. 146, 368–376

54. Sparkes, I. A., Runions, J., Kearns, A., and Hawes, C. (2006) Rapid, transient expression of fluorescent fusion proteins in tobacco plants and generation of stably transformed plants. Nat Protoc. 1, 2019–2025

