## Supplemental information for "The cytosolic *Arabidopsis thaliana* cysteine desulfurase ABA3 delivers sulfur to the sulfurtransferase STR18"

### SUPPORTING INFORMATION

**Table S1.** Primers used for cloning and site-directed mutagenesis experiments.

**Figure S1.** Biochemical properties of the CD-Rhd C466S variant.

**Figure S2.** STR18 exhibits Cys89-dependent TST activity.

**Figure S3.** STR18 does not use L-Cys as a sulfur donor.

**Figure S4.** Electrospray ionization mass spectrometry analysis of STR18.

**Figure S5.** Electrospray ionization mass spectrometry analysis of the STR18 C47S variant.

**Figure S6.** Electrospray ionization mass spectrometry analysis of the STR18 C89S variant.

**Figure S7.** STR16 catalyzes the oxidation of roGFP2 via trans-persulfidation.

**Table S1. Primers used for cloning and site-directed mutagenesis experiments.**
The *Nde*I, *Bam*HI, *Kpn*I and *Mlu*I restriction sites used for cloning are underlined in the primers. The mutagenic codons are in bold.

| **Name** | **Sequence** |
|---|---|
| AtSTR18 for | 5’ CCCCCCCCCATATGGCTTCTCAATCAATCTCCTCCAGC 3’ |
| AtSTR18 rev | 5’ CCCCGGATCCTTAATTAGCAGATGGCTCCTC 3’ |
| AtSTR18 C47S for | 5’ TTTAGGAGAGGCCAT**TCT**GAGGCAGCTAAGATC 3’ |
| AtSTR18 C47S rev | 5’ GATCTTAGCTGCCTC**AGA**ATGGCCTCTCCTAAA 3’ |
| AtSTR18 C89S for | 5’ GATATCCTTGTGGGT**TCT**CAGAGTGGAGCCAGA 3’ |
| AtSTR18 C89S rev | 5’ TCTGGCTCCACTCTG**AGA**ACCCACAAGGATATC 3’ |
| AtABA3 for | 5’ CCCCCCCCCATATGGAAGCATTTCTTAAGGAATTC 3’ |
| AtABA3 rev | 5’ CCCCGGATCCTTATTCAATATCTGGATTAACTTC 3’ |
| AtSTR16 for | 5’ CCCCCCCCCATATGGCGGAGGAGAGCAGAGTC 3’ |
| ASTR16 rev | 5’ CCCCGGATCCTTAAGCCTTTGTAGGAAGGCC 3’ |
| AtSTR16 C80S for | 5’ AACATCATTGTTGGC**AGC**CAGAGCGGTGGTAGA 3’ |
| ASTR16 C80S rev | 5’ TCTACCACCGCTCTG**GCT**GCCAACAATGATGTT 3’ |
| AtACL1 cLuc for | 5’ CCCCCCGGTACCATGGCGAGGAAGAAGATC 3’ |
| AtACL1 cLuc rev | 5’ CCCCCCGGATCCTCATGCTGCTGCTGTGAT 3’ |
| ABA3 nLuc/cLuc for | 5’ CCCCCCGGTACCATGGAAGCATTTCTT 3’ |
| ABA3 nluc rev | 5’ CCCCCCACGCGTGTTCAATATCTGGATT 3’ |
| ABA3 cluc rev | 5’ CCCCCCGGATCCTCATTCAATATCTGGATT 3’ |
| STR18 nLuc/cLuc for | 5’ CCCCCCGGTACCATGTCTCAATCAATC 3’ |
| STR18 nluc rev | 5’ CCCCCCACGCGTGATTAGCAGATGGCTC 3’ |
| STR18 cluc rev | 5’ CCCCCC**GGATCC**TCAATTAGCAGATGGCTC 3’ |
| CD-Rhd C466S for | 5’ CAGGTCATCTTCGTG**TCC**CGCAGCGGCCGGCGC 3’ |
| CD-Rhd C466S rev | 5’ GCGCCGGCCGCTGCG**GGA**CACGAAGATGACCTG 3’ |

**Figure S1**

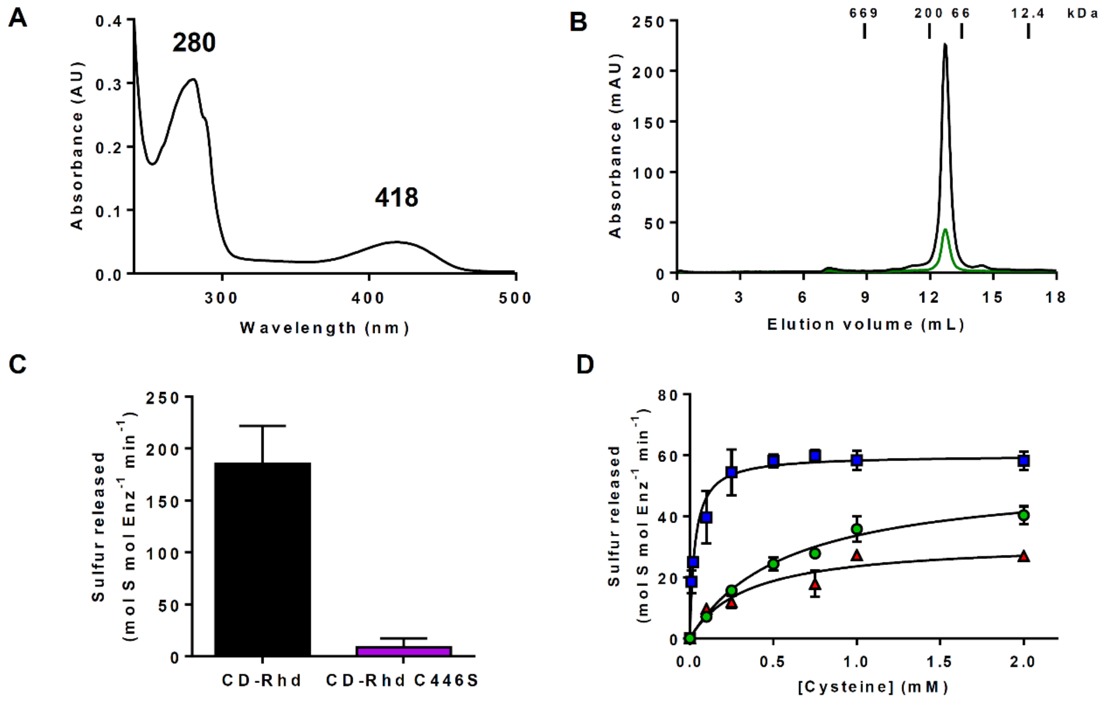


**Figure S1. Biochemical properties of the CD-Rhd C466S variant.** A. UV-visible absorption spectrum of the purified N-terminal His-tagged recombinant CD-Rhd C466S recorded in a 30 mM Tris-HCl pH 8.0 buffer. B. Analytical gel filtration (Superdex S200 10/300 column, GE Healthcare) of His-tagged recombinant CD-Rhd C446S (100 µg). The presence of the polypeptide and of the PLP cofactor have been detected by measuring the absorbance at 280 nm (dark line) and 418 nm (grey line), respectively. The apparent molecular weight of CD-Rhd was estimated from the separation of the indicated standards. C. Comparison of thiosulfate sulfurtransferase activity between CD-Rhd and its C466S variant. Reactions were performed in the presence of 100 nM CD-Rhd, 5 mM thiosulfate and 5 mM β-mercaptoethanol. The data are represented as mean ± SD of three independent experiments. D. Steady-state kinetic parameters of the cysteine desulfurase activity. Reactions were performed in the presence of 100 nM CD-Rhd C466S, increasing concentrations of L-cysteine (0 to 2 mM) and in the presence of various reductants, either 5 mM of DTT (blue squares), or 5 mM GSH (green circles) or 5 mM β-mercaptoethanol (red triangles). The data are represented as mean ± SD of three independent experiments.

**Figure S2**

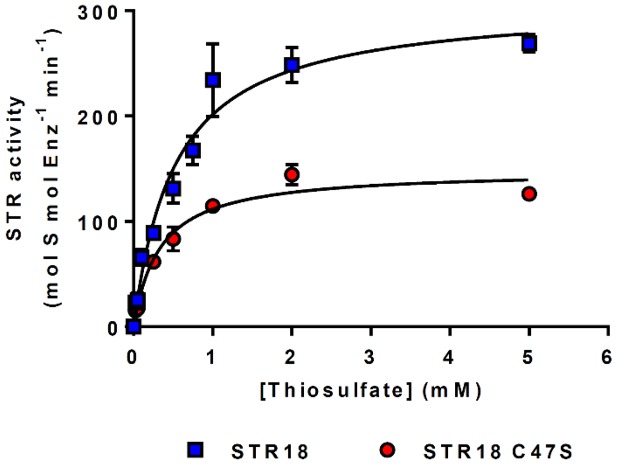


**Figure S2. The TST activity of STR18 requires Cys89.** The TST activity of STR18 (blue squares), STR18 C47S (red squares) was measured under steady-state conditions using the lead acetate method in the presence of 100 nM STR18, 5 mM β-mercaptoethanol and increasing concentrations of thiosulfate (0 to 5 mM). The data are represented as mean ± SD of three independent experiments.

**Figure S3**

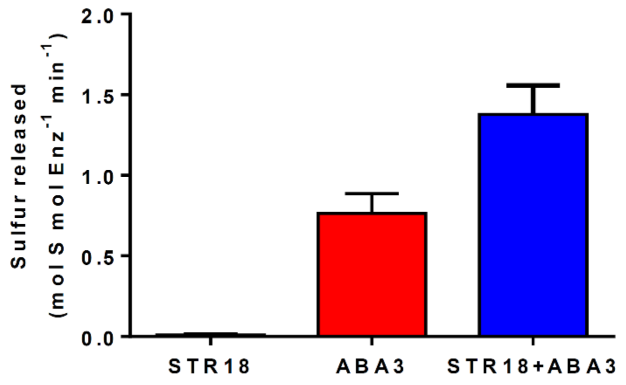


**Figure S3. STR18 does not use cysteine as a sulfur donor.** CD activity was measured in the presence of 250 µM L-cysteine and 5 mM DTT with either 5 µM STR18 or 1 µM ABA3 alone and with 5 µM STR18 as described in the 'Experimental procedures" section. The data are represented as mean ± SD of three independent experiments.

**Figure S4**

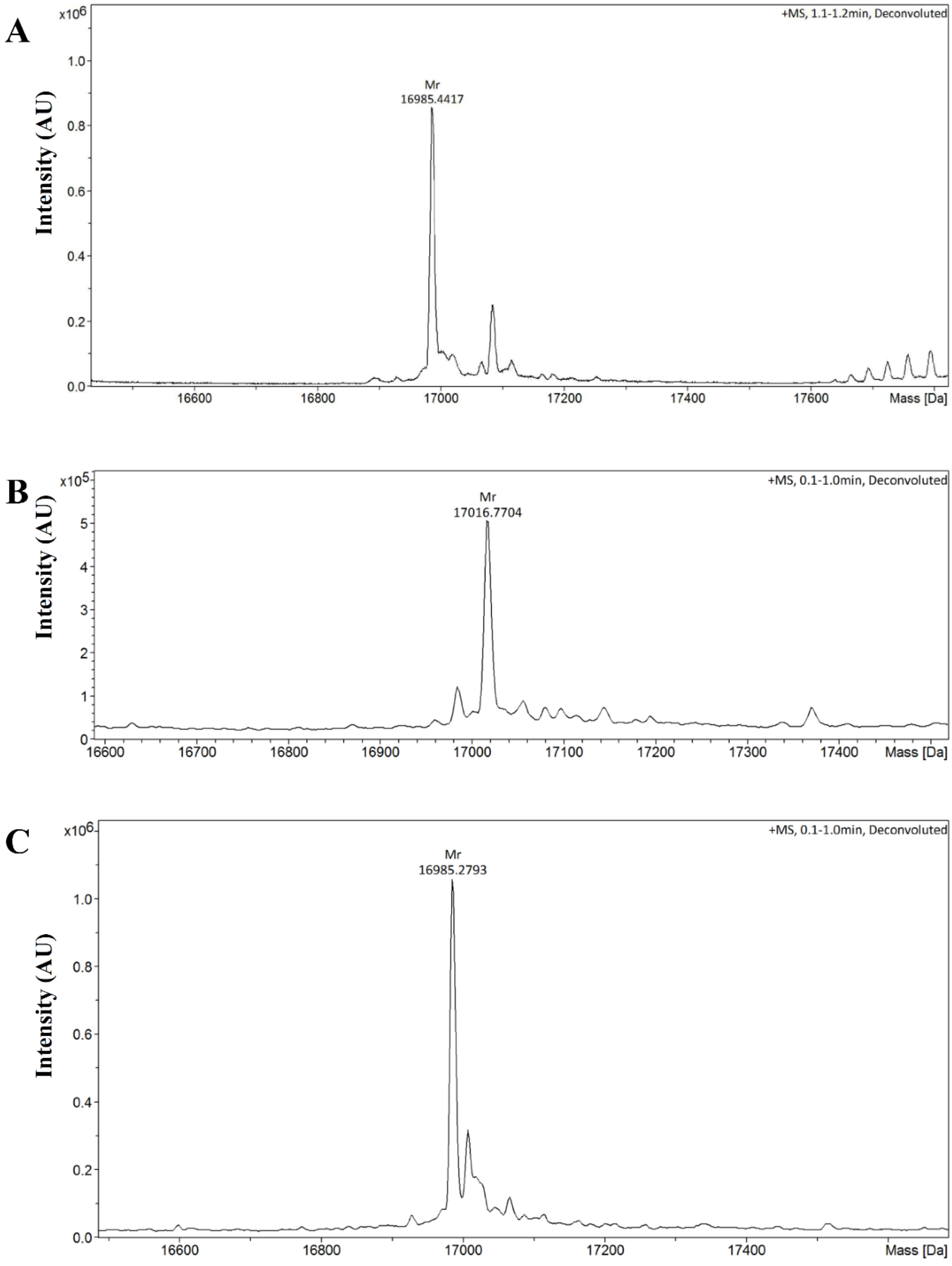


**Figure S4. Electrospray ionization mass spectrometry analysis of STR18.** Deconvoluted mass spectra of STR18 determined for a reduced protein (A), a reduced protein incubated with L-cysteine and ABA3 before (B) or after subsequent treatment with DTT (C) as described in the "Experimental procedures" section.

**Figure S5**

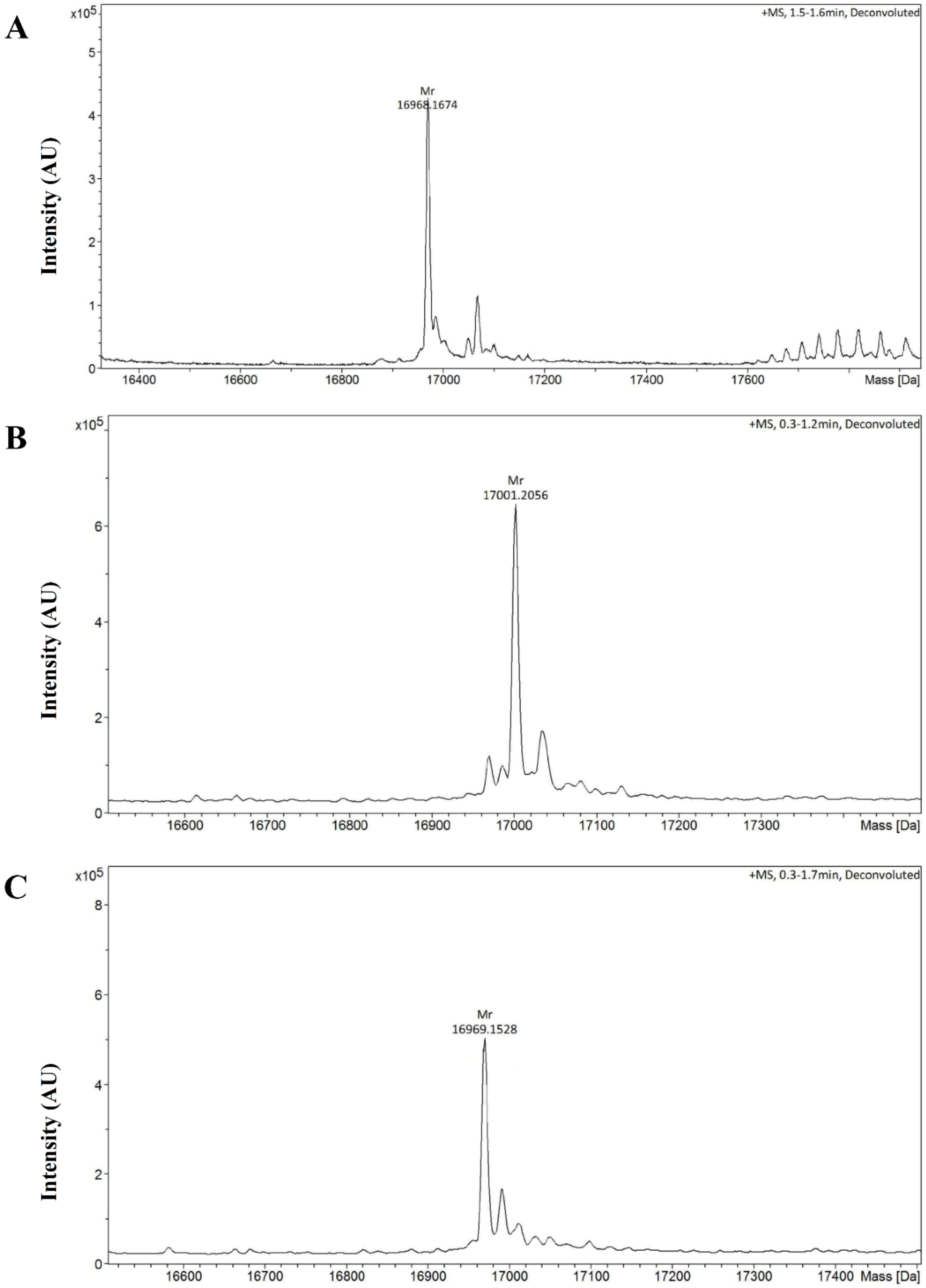


**Figure S5. Electrospray ionization mass spectrometry analysis of the STR18 C47S variant.** Deconvoluted mass spectra of STR18 C47S determined for a reduced protein (A), a reduced protein incubated with L-cysteine and ABA3 before (B) or after subsequent treatment with DTT (C) as described in the “Experimental procedures” section

**Figure S6**

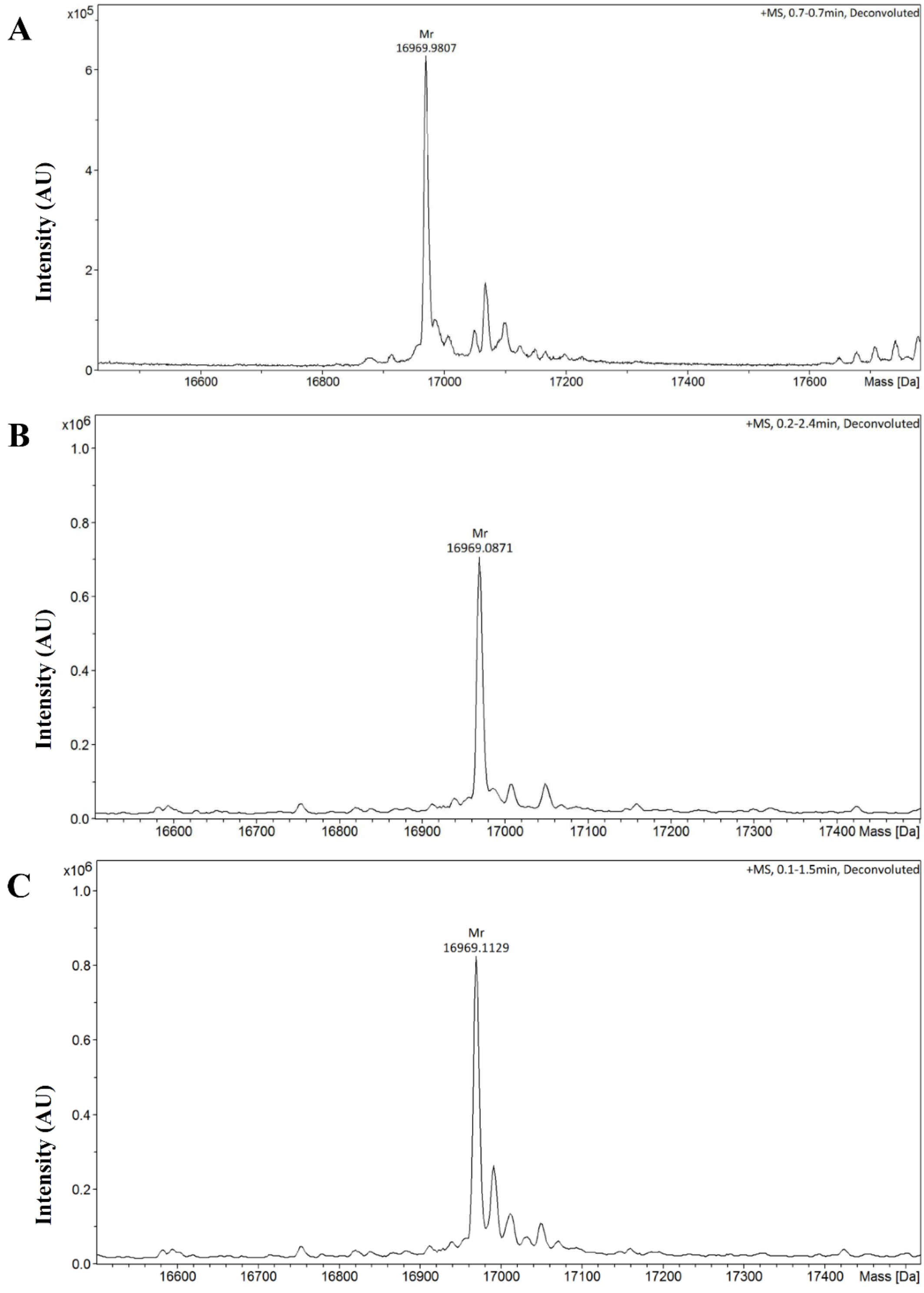


**Figure S6. Electrospray ionization mass spectrometry analysis of the STR18 C89S variant.** Deconvoluted mass spectra of STR18 C89S determined for a reduced protein (A), a reduced protein incubated with L-cysteine and ABA3 before (B) or after subsequent treatment with DTT (C) as described in the "Experimental procedures" section.

**Figure S7**

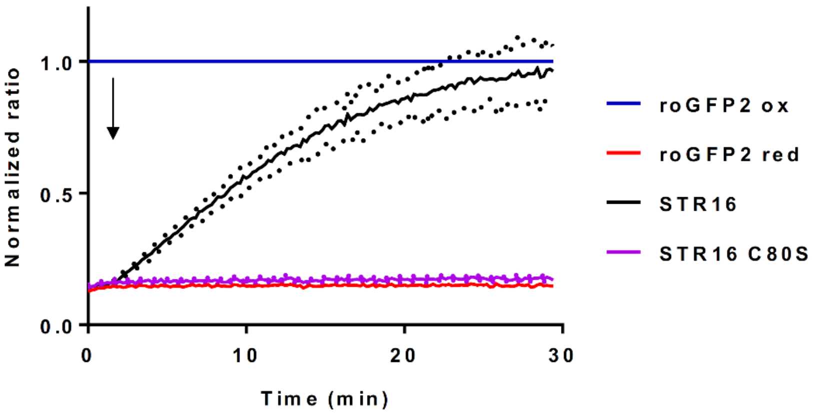


**Figure S7. STR16 catalyzes the oxidation of roGFP2 via trans-persulfidation.** Persulfide-dependent oxidation kinetics of 1 µM roGFP2 in the presence of 5 µM STR16 and 5 mM thiosulfate. STR16 C80S variant was used as control of inactive STR16. The arrows indicate the addition of thiosulfate after 2 min. The fully reduced or oxidized roGFP2 used as references were obtained after incubation with 10 mM DTT or $H_2O_2$ respectively. The ratio of 400/480 nm was normalized to the respective value of maximal roGFP2 oxidation by $H_2O_2$. The data are represented as mean ± SD of three independent experiments.
